# Engineering combinatorial and dynamic decoders using synthetic immediate-early genes

**DOI:** 10.1101/2019.12.17.880179

**Authors:** Pavithran T. Ravindran, Maxwell Z. Wilson, Siddhartha G. Jena, Jared E. Toettcher

## Abstract

For tissues to grow and function properly, cells must coordinate actions such as proliferation, differentiation and apoptosis. This coordination is achieved in part by the activation of intracellular signaling pathways that trigger the expression of context-specific target genes. While the function of these natural circuits has been actively studied, synthetic biology provides additional powerful tools for deconstructing, repurposing, and designing novel signal-decoding circuits. Here we report the construction of synthetic immediate-early genes (synIEGs), target genes of the Erk signaling pathway that implement complex, user-defined regulation and can be monitored through the use of live-cell biosensors to track transcription and translation. We demonstrate the power and flexibility of this approach by confirming Erk duration-sensing by the *FOS* immediate-early gene, elucidating how the *BTG2* gene is regulated by transcriptional activation and translational repression after growth-factor stimulation, and by designing a synthetic immediate-early gene that responds with AND-gate logic to the combined presence of growth factor and DNA damage stimuli. Our work paves the way to defining the molecular circuits that link signaling pathways to specific target genes, highlighting an important role for post-transcriptional regulation in signal decoding that may be masked by analyses of RNA abundance alone.

## Introduction

In mammals, relatively few intracellular pathways integrate information from a huge range of sources, including neighboring cells and the physical environment. The resulting activity of signaling pathways can have many consequences, but chief among these is the induction of target genes that constitute a cell’s decision to proliferate, differentiate, or adopt an altered functional state. Efforts to systematically map signaling responses have revealed that a single external stimulus often activates many intracellular pathways, and that each pathway’s activity state can vary dynamically over time (*1–4*). These observations suggest that cells might use both combinatorial strategies (e.g., gene expression triggered only in response to pathways A and B) and dynamic strategies (e.g., gene expression triggered by sustained activity in pathway A) to connect the cell’s overall signaling state to particular responses. Indeed, a growing number of cell fates are thought to be selectively triggered by certain signaling dynamics, thereby functioning as analog filters (**Figure 1A**) (*5–7*), whereas others may act as digital logic gates by responding only to certain pathway combinations (**Figure 1B**) (*8*). Yet our understanding of how combinatorial and dynamic decoding are achieved is still limited.

**Fig 1.**
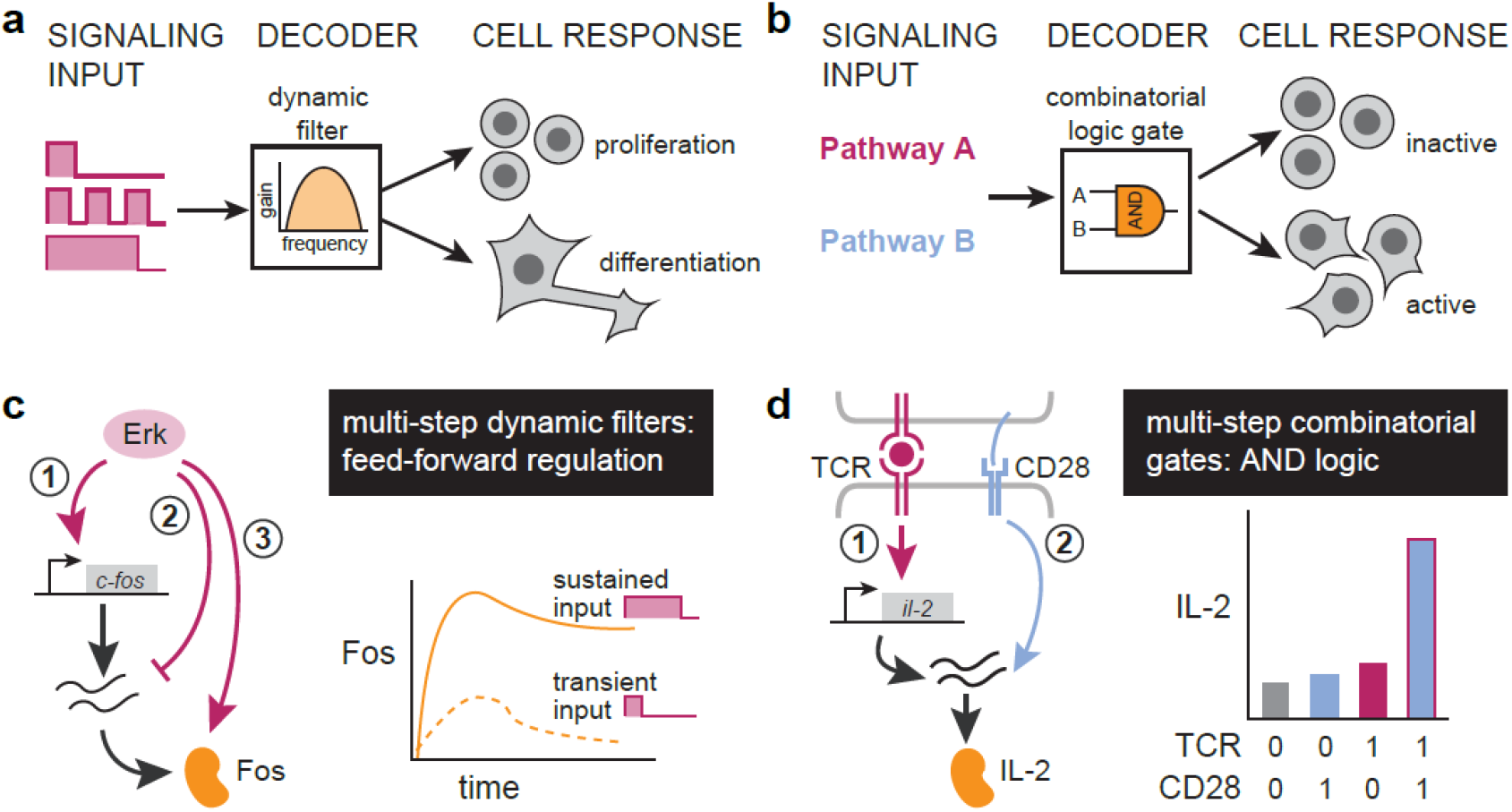
Cells use multistep regulation to interpret dynamic and combinatorial signaling inputs. (**A**) Target gene induction may depend on the dynamics of signaling pathway activation, such as the duration, frequency or area-under-the-curve of pathway activity. In such cases, the signal-decoding circuitry may be thought of as a dynamic filter. (**B**) Target gene induction may also depend on the combination of pathways that are activated, such that signal decoding may be thought of as implementing a logic gate. (**c**) The induction of Fos protein is a canonical example of dynamic decoding, where sustained but not transient pulse of Erk results in protein accumulation. Erk-mediated regulation of *FOS* transcription, *fos* mRNA stability and Fos protein stability is thought to mediate this response. (**D**) IL-2 induction by T cell stimulation and co-stimulation is thought to occur *via* combinatorial control. Neither TCR nor CD28 alone are sufficient for protein output but the two together allow for accumulation of IL-2 protein through a poorly-characterized response circuit.

A central challenge is that the relationship between signaling and target gene activation is complex, with multiple regulatory links acting at different steps along the central dogma, an architecture we will call “multi-step regulation.” One canonical example of multi-step regulation is found in *FOS* gene induction by the Ras/Erk pathway. Erk signaling first triggers transcription of *fos* mRNA; then, within 30-60 min, *fos* mRNA is degraded through the Erk-induced expression of Zfp36; and finally, the Fos protein is stabilized by Erk phosphorylation (*9–12*). Together, these interactions are thought to form a circuit that selectively responds only when the duration of Erk activity is above a threshold (**Figure 1C**). Multi-step regulation can also provide combinatorial control when sequential steps in gene expression are gated by distinct signaling pathways. For instance, in T cells, engagement of the T cell receptor (TCR) leads to *il-2* transcription, but maximal IL-2 secretion requires CD28-dependent signaling acting at post-transcriptional steps that are still poorly defined (**Figure 1D**), resulting in AND-gate logic where both TCR and CD28 engagement are required for a strong cytokine response (*13*). In both cases, the complexity of multiple nested regulatory links has made it challenging to define the essential set of interactions implement a specific filtering or gating function. A more complete understanding of multi-step regulation would also enable the design of synthetic decoding modules: gene circuits that selectively respond to novel stimulus combinations or dynamics.

Motivated by these challenges, we set out to establish a general framework for constructing synthetic, signaling-responsive target genes that can be used to implement user-defined, multistep regulatory interactions. We focused specifically on creating and characterizing synthetic immediate-early genes (SynIEGs), a class of fast-responding genes that are induced by a variety of stimuli including the Ras/Erk signaling pathway. We found that SynIEGs faithfully recapitulate the dynamics of immediate-early gene induction: SynIEG transcription kinetics closely match their endogenous counterparts, and a *FOS*-based SynIEG exhibits dynamic filtering of Erk signaling inputs. We also use the SynIEG platform to define additional regulatory links, revealing an essential role for the *BTG2* 3’ UTR and the micro-RNA *miR-21* in suppressing Btg2 protein translation. Finally, we use regulatory elements from the *FOS* and *BTG2* genomic loci to engineer a SynIEG with a novel decoding function: an AND gate that selectively responds only to the combination of growth factor and DNA damage stimuli. Synthetic signaling-responsive target genes thus enable a quantitative, systems-level understanding of the interface between signaling pathways and gene expression, opening the door to engineering novel pathway decoders for controlling complex cell fates.

## Results

### Developing a synIEG platform for monitoring synthetic target gene induction

Our strategy for constructing synthetic target genes relied on meeting two complementary goals. First, a SynIEG must be able to easily implement different forms of regulation at the mRNA or protein level. Each SynIEG thus combines a signaling-responsive promoter, 5’ and 3’ mRNA regulatory sequences, and protein-coding sequences into a vector that can be targeted for genomic integration (**Figure 2A**). Second, a quantitative understanding of signal decoding requires the ability to monitor mRNA- and protein-level responses over time in each individual SynIEG-expressing cell. Building off of a strategy we recently developed for endogenous target genes (*14*), we included a YFP-24xMS2 tag in each SynIEG. In this system, instantaneous transcription can be visualized as a bright nuclear spot when the nascent MS2 RNA loops are bound to the fluorescent RNA-binding protein MCP-mCherry, and protein accumulation can be monitored by YFP fluorescence. As a first test case, we combined a 2-kb region upstream of the *FOS* transcriptional start site that contains its promoter and canonical upstream regulatory elements (*15, 16*), the *FOS* 5’UTR, the coding sequence for monomeric super-folder YFP (msfYFP), 24xMS2 RNA stem-loops, and the *TUBA1B* 3’UTR in a single lentiviral vector, which we named *fos-tubulin* for its 5’ and 3’ elements, respectively (**Figure 2B**), and introduced it into a clonal NIH3T3 cell line already expressing MCP-mCherry and H2B-iRFP as a nuclear marker (the “chassis” cell line; see Methods) (*14*).

**Fig 2.**
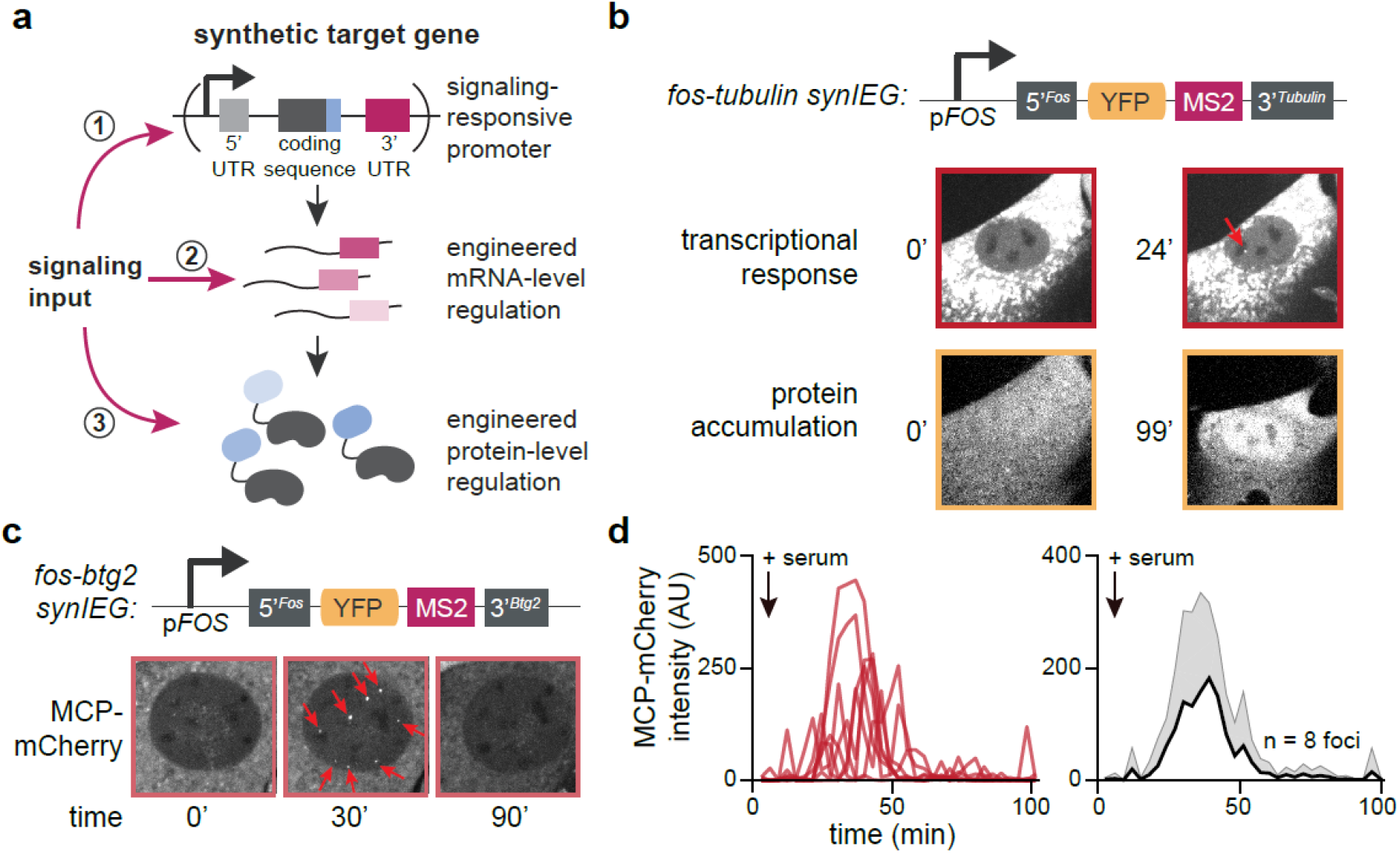
Development of synthetic Ras/Erk target genes that recapitulate endogenous transcriptional kinetics. (**A**) Schematic overview of synthetic target genes incorporating multistep regulation. Signaling inputs can act at the transcriptional level through promoter/enhancer regulation (1), the mRNA level through UTR-based regulation (2), and at the protein level through signaling-responsive domains or linear motifs (3). (**B**) Design of synthetic immediate-early genes (synIEGs). The Erk responsive *FOS* promoter (P_Fos_) drives the expression, which allows for the visualization of transcription and translation in live cells, allowing one to study the information flow from signaling input to protein accumulation. mRNA UTR elements and protein degrons can be added to modulate mRNA and protein stability. (**C**) Representative images of synIEG contain *fos* 5’ UTR and *tubulin* 3’ UTR before and after serum stimulation for both transcription and translation. Transcriptional focus is denoted with a red arrow. (**D**) Images of NIH 3T3 cells after induction of transcription of synIEG contain *fos* 5’ UTR and *btg2* 3’ UTR at multiple time-points after serum stimulation. Transcriptional foci are denoted using red arrows. Quantification of transcriptional foci in (d), n=8 foci showing (left) individual traces and (right) mean + SD.

We first tested whether expression of the *fos-tubulin* synthetic immediate-early gene (SynIEG) could be induced by signaling stimuli after either lentiviral transduction or transient transfection, but found that neither strategy was promising. Transient transfection with a *fos-tubulin* plasmid drove YFP expression even in the absence of IEG-activating stimuli such as serum (**Figure S1A-B**). Conversely, cells transduced with a *fos-tubulin* lentiviral vector failed to induce YFP expression even after serum stimulation (**Figure S1C**). Based on prior reports that lentiviral targeting constructs can be silenced (*17*), we reasoned that treatment with the histone deacetylase inhibitor trichostatin A (TSA) might restore serum responsiveness to a lentiviral *fos-tubulin* cell line. Indeed, cells pre-treated with TSA for 12 h exhibited an increase in YFP fluorescence upon serum stimulation, whereas cells treated with either TSA or serum alone showed no change in YFP levels (**Figure S1C-D**). While these data indicate that lentiviral-transduced SynIEGs can be reactivated from a silent state, the large-scale chromatin remodeling caused by TSA treatment made this an undesirable strategy for implementing SynIEGs.

As an alternative method of synthetic gene delivery, we tested integration using the PiggyBAC transposase, which randomly inserts DNA sequences flanked by ~300-bp targeting sequences into the host genome (*18*). We co-transfected “chassis” NIH3T3 cells with plasmids encoding the PiggyBAC transposase enzyme and the *fos-tubulin* SynIEG flanked by PiggyBAC transposable elements, and sorted clonal cell lines that stably integrated the SynIEG (**Figure 2B**). We observed that *fos-tubulin* cells stimulated with serum exhibited bright MCP-labeled transcriptional foci and increased YFP fluorescence over time, consistent with an Erk-stimulated IEG response (**Figure 2B, Figure S1E-F**). In contrast to lentiviral transduction, silencing was not observed and cells maintained serum-responsiveness even months after cell line generation. These results confirm that PiggyBAC transposase-based delivery enables the stable integration of complex, signaling-responsive synthetic target genes for interrogating mRNA- and protein-level responses.

How well do SynIEGs recapitulate the dynamics of endogenous immediate-early gene activation? To address this question we monitored transcription at individual genomic loci using the MS2/MCP system built into each SynIEG. In this system, the transcription rate of each genomic integration site can be tracked over time based on the intensity of individual fluorescent MCP foci in the nucleus. We generated a clonal cell line harboring multiple integrations of a *fos-btg2* SynIEG (containing the *FOS* 5’ regulatory sequence and *BTG2* 3’ UTR), based on our prior work demonstrating that *BTG2*’s long 3’ UTR produces exceptionally bright transcriptional foci (*14*). Upon serum stimulation, *fos-btg2* cells exhibited multiple bright transcriptional foci, each corresponding to transcription from distinct PiggyBAC integration sites (**Figure 2C**). Focus intensity reached a maximum intensity roughly 30 minutes after stimulation and then adapted back to baseline within 90 minutes (**Movie S1; Figure 2D**). Transcriptional kinetics were strikingly similar between distinct foci within the same cell, as well as between cells in the same cell line (**Movie S1; Figure 2D**).

We next set out to compare the kinetics of serum-stimulated SynIEG transcription with their endogenous IEG counterparts. We had previously established derivatives of the “chassis” NIH3T3 cell line where MS2 stem-loops were integrated at the endogenous *FOS* and *BTG2* loci, providing an ideal testbed for comparison (*14*). We stimulated *fos-btg2* SynIEG cells and these endogenously-tagged *FOS* and *BTG2* cell lines with 10% serum (**Figure S2A**), and quantified both the kinetics of transcriptional activation and subsequent adaptation to the baseline state (**Figure S2B**). We found that activation and adaptation in *fos-btg2* SynIEG cells was strikingly similar to those of the endogenous *FOS* genomic locus, suggesting that *FOS* bursting kinetics are solely controlled by the proximal *FOS* enhancer-promoter sequence in our synthetic gene construct (**Figure S2C**). SynIEGs also captured many prior observations about immediate-early gene regulation (*14*): the transient pulse of SynIEG transcription was converted to a sustained transcriptional response by co-treatment with the protein synthesis inhibitor cycloheximide (**Figure S2D-E**), and serum-induced transcription was immediately blocked by pharmacological inhibition of the MAPK pathway (**Figure S2F**). Taken together, these data indicate that transposase-integrated SynIEGs are highly sensitive to upstream MAPK pathway signaling and suggest that SynIEGs quantitatively recapitulate the dynamics of gene expression of their endogenous IEG counterparts.

### SynIEGs elucidate dynamic and combinatorial decoding by *FOS* and *BTG2*

We next tested whether SynIEGs could be used to define the regulatory steps that enable target genes to selectively respond to certain combinations or dynamics of upstream signaling. We focused on the *FOS* and *BTG2* genes, two immediate-early genes that exhibit distinct profiles of signaling-dependent protein expression. *FOS* is a canonical example of dynamic decoding as it is highly induced by sustained, but not transient, activation of the Ras/MAPK pathway (**Figure 3A**) (*11, 12*). In contrast, *BTG2* induction appears to depend on the specific stimulus received: Btg2 protein accumulates to high levels upon DNA damage but not growth factor stimulation, even though *btg2* mRNA is strongly induced in both cases (*14*). We thus set out to construct variants of *FOS* and *BTG2*-based SynIEGs to recapitulate and further define their regulatory architecture.

**Fig 3.**
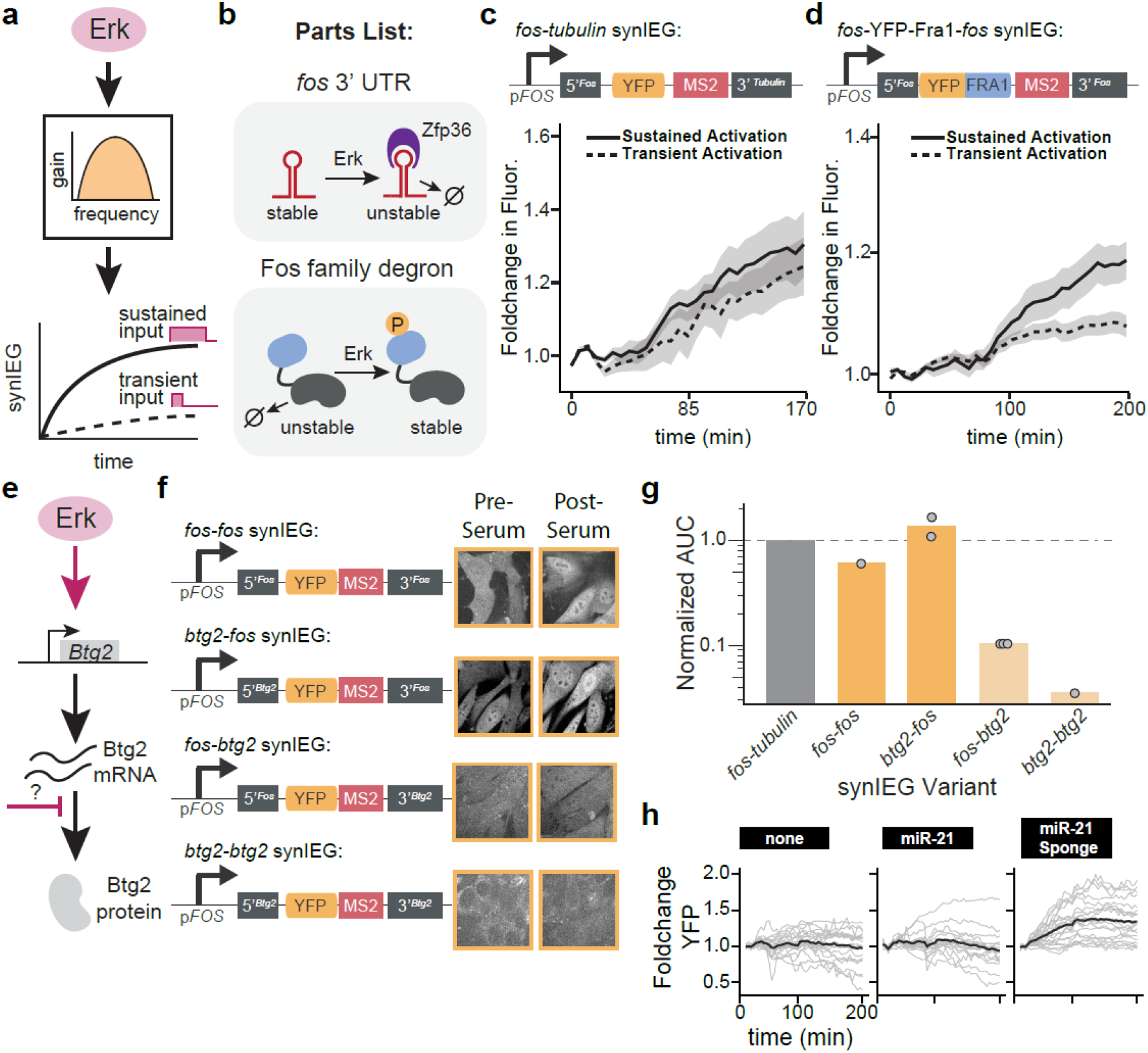
synIEGs can be used to implement dynamic and combinatorial decoding circuits. (**A**) Schematic overview of a dynamic decoding circuit based on *FOS* regulatory logic. Erk activates a SynIEG (middle box), enabling sustained but not transient induction of target protein levels. (**B**) Parts list of Erk-dependent regulation for the *FOS* immediate-early gene. The *fos* 3’ UTR is subject to degradation by a second Erk-induced immediate-early gene, Zfp36 (upper panel). Conversely, Fos protein levels are stabilized by Erk phosphorylation of a degron sequence shared by Fos family members (lower panel). (**C-D**) Quantification of YFP induction as a function of stimulus duration for the *fos-tubulin* SynIEG (in **C**) and the *fos*-Fra1^deg^-*fos* SynIEG implementing Fos-specific mRNA- and protein-level regulation (in **D**). The fold-change in YFP fluorescence was quantified from NIH 3T3s that also expressed the iLID-OptoSOS system and were stimulated with either transient (30 min) or sustained (90 min) blue light. All curves indicate the mean ± S.E.M for n=10 cells (panel **C**, sustained), n=6 cells (panel **C**, transient), n=22 cells (panel **D**, sustained), and n=23 cells (panel **D**, transient). (**E**) Schematic of combinatorial control over *BTG2* induction: Erk stimulation induces *BTG2* transcription but not protein accumulation, whereas DNA damage induces both. The mechanism underlying this difference is unknown. (**F**) Representative images of YFP levels in NIH 3T3 clonal lines expressing various SynIEGs and stimulated with 10% serum. Images show YFP levels just before and 3 h after serum stimulation. (**G**) Quantification of the area-under-the-curve (AUC) of YFP induction after serum stimulation for the clonal SynIEG cell lines shown in **F**, as well as for *fos-tubulin* as an additional control (see Methods for quantification details). Each point represents the average of 20-30 cells from an independent experiment. (**H**) Quantification of YFP fluorescence in NIH3T3 cells harboring the *fos-btg2* synIEG and stimulated with 10% serum after being transduced with nothing (n=20 cells), *miR-21* (n=17 cells) or the *miR-21* sponge (n= 21 cells). Single-cell traces are shown in gray, with the mean response shown as a black line.

Dynamic decoding of *FOS* is thought to depend on two forms of post-transcriptional regulation – Erk-dependent destabilization of the *FOS* 3’ UTR (*19*) and stabilization of Fos protein (*11*) – although additional forms of regulation have not been ruled out. We thus tested whether this minimal set of regulatory elements would indeed be capable of dynamic filtering. We combined the *FOS* promoter, *FOS* 3’UTR, and a well-characterized Erk-responsive degron from the FOS family member Fra1 (*20*) to build a SynIEG termed *fos*-Fra1^deg^-*fos* (**Figure 3B**). The *fos-tubulin* SynIEG was used as a control lacking both forms of post-transcriptional regulation. We introduced both SynIEGs into “chassis” NIH3T3 cells that were also sorted to express a blue light sensitive OptoSOS system (iLID-OptoSOS) (16, 17), enabling us to precisely control the dynamics of Ras/Erk pathway activity by varying the duration of illumination (**Figure S3A**). In the absence of blue light, SSPB-SOScat is cytosolic and inactive, whereas blue light stimulation induces SSPB-SOScat membrane localization (**Figure S3B**) and activates Ras/Erk signaling (**Figure S3C-E**).

We incubated OptoSOS-SynIEG cells in serum-free media for 6 h, applied either a 30 min pulse of light (‘transient’) or continuous illumination (‘sustained’), and monitored YFP induction over time (**Figure 3C-D**). For *fos-tubulin* cells, both light stimuli triggered similar increases in YFP fluorescence over time (**Figure 3C**), likely because both sustained and transient Erk-activating stimuli cause an identical, 30 min pulse of transcription (**Figure 2**) (*14*). However, we found that in the *fos*-Fra1^deg^-*fos* clonal cell line, sustained light drove higher protein accumulation than a 20 min light pulse (**Figure 3D**). It is important to note that stimulation led to a relatively low overall change in SynIEG protein expression: 40% in the case of *fos-tubulin* and 20% in the case of *fos*-Fra1^deg^-*fos* cells over a 3 h timecourse (**Figure 3C-D**). This observation is likely to arise from a high residual pool of stable YFP protein that is retained from continuous growth conditions prior to the start of the experiment, a hypothesis we further tested in **Figure 4** below. Nevertheless, the observation of dynamic selectivity allows us to conclude that the two post-transcriptional regulatory connections contained in the *fos*-Fra1^deg^-*fos* SynIEG (Erk-dependent protein stabilization and mRNA degradation) are sufficient to confer dynamic selectivity for sustained Erk stimuli.

**Fig 4.**
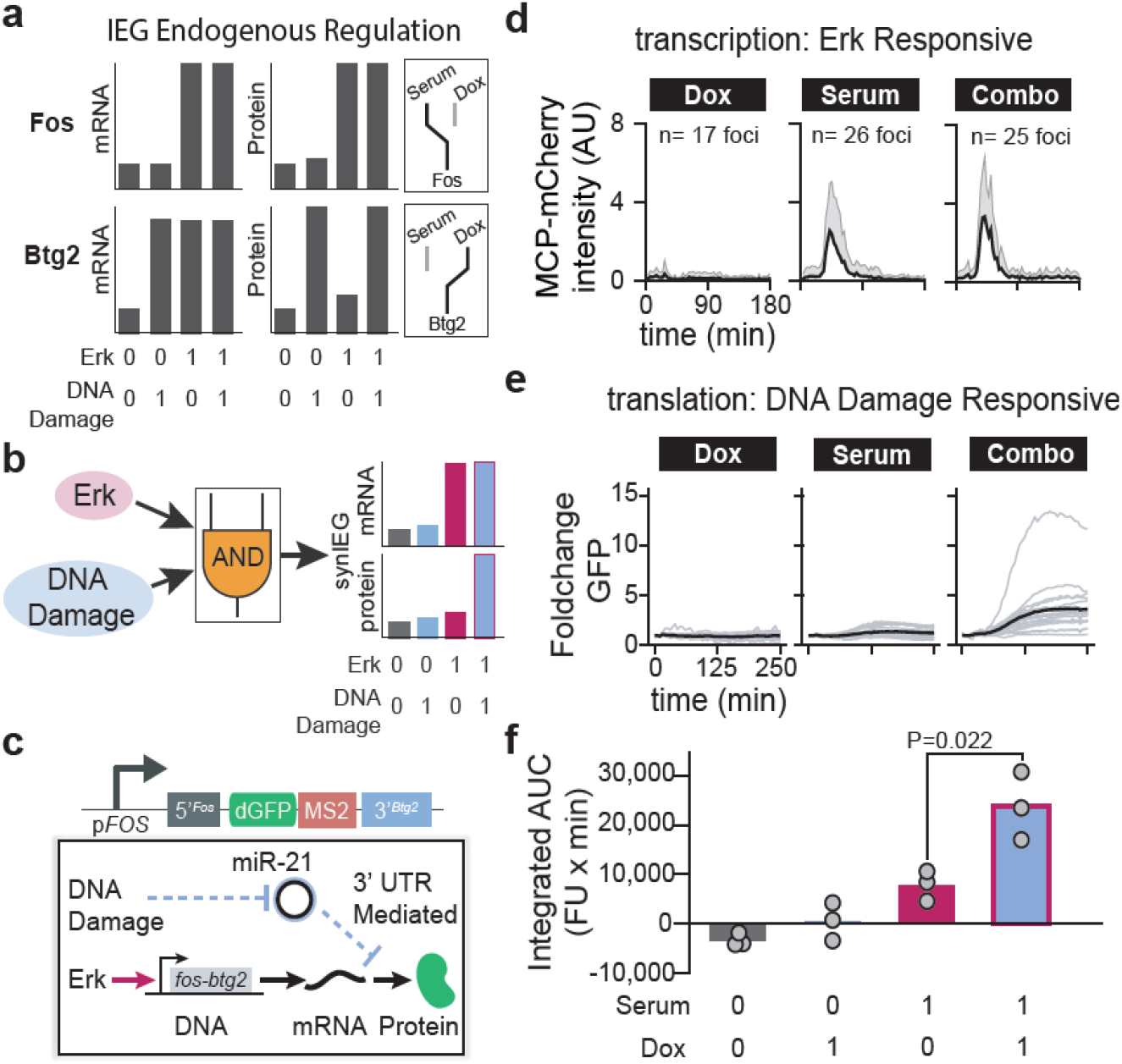
A SynIEG-based AND gate triggers target gene expression in response to Erk and DNA damage. (**A**) Cartoon schematic of prior knowledge of Fos and Btg2 regulation in response to growth factor stimulation (Serum) and DNA damage (doxorubicin; Dox). Fos transcription and protein accumulation are both triggered by serum stimulation irrespective of DNA damage (upper panels). In contrast, Btg2 transcription is induced by both DNA damage and serum stimulation, yet only DNA damage triggers Btg2 protein accumulation. (**B**) Schematic of a synthetic AND gate responding to both serum stimulation and DNA damage. By combining a *FOS*-like transcriptional response with a *BTG2*-like protein response, target gene induction would be triggered only when both stimuli are supplied. (**C**) Experimental implementation of the AND-gate SynIEG. Transcription of the *fos-btg2* SynIEG is induced only by serum stimulation, not DNA damage, due to its use of the *FOS* 3’UTR. However, DNA damage is required for protein accumulation through the use of the *BTG2* 3’UTR. (**D**) Transcriptional response of the *fos-btg2* SynIEG. The brightness of individual nuclear MCP-mCherry foci was quantified (mean + S.D.) in cells treated with doxorubicin (n=17 foci), serum (n = 26 foci), or their combination (n=25 foci). (**E**) GFP induction from the *fos-btg2* SynIEG. Cytosolic dGFP levels were quantified from cells stimulated with doxorubicin, serum, or their combination. Single-cell traces are shown in gray, with the mean response shown as a black line. (**F**) Quantification of the area-under-the-curve (AUC) of GFP induction for the clonal SynIEG cell lines shown in **D-E** after stimulation with serum, doxorubicin, or their combination (see Methods for quantification details). Each point represents the mean of at least 20 cells in a single experiment, with each bar representing the mean of three independent replicate experiments. The p-value was computed using the Student’s t-test between serum-only and serum + dox conditions.

We next turned our attention to *BTG2*, whose regulation by various upstream signals is still poorly understood (*21*). *BTG2* is transcriptionally induced by many cellular inputs, including DNA damage (*22, 23*), growth factor signaling (*14*), and stress signaling (*24*). Strikingly, we previously saw that Btg2 protein levels were unchanged after growth factor stimulation despite strong transcriptional induction (**Figure 3E**) (*14*). Our first goal was to determine whether the block to protein accumulation occurred at the mRNA level (*e.g*. by regulating mRNA degradation or blocking translation) or at the protein level (*e.g*. by regulating protein degradation). To determine whether mRNA-level regulation is sufficient, we first constructed a series of SynIEG variants that lacked any Btg2 protein sequence but harbored various combinations of *BTG2* and *FOS* UTR sequences (termed the *btg2-btg2, btg2-fos, fos-btg2*, and *fos-fos* SynIEGs). We derived clonal cell lines for each SynIEG, starved each for 4-6 hours, and then stimulated with serum to monitor YFP induction, using identically-treated *fos-tubulin* SynIEG cells as controls (**Figure S4**). We found that all SynIEGs containing the *BTG2* 3’ UTR failed to accumulate YFP in response to serum stimulation (**Figure 3F-G**), whereas all other SynIEGs exhibited similar levels of YFP accumulation. Together, these data demonstrate that RNA-level regulation is sufficient to explain Btg2’s paradoxical response of high serum-induced expression with no protein accumulation, and implicates regulation contained within the *BTG2* 3’ UTR.

Prior reports indicate that over a dozen microRNAs may bind to the *BTG2* 3’ UTR and regulate *btg2* mRNA stability or protein expression in various cell types and cancers (*25–29*). We focused on *miR-21* as a candidate regulator because it is upregulated after growth factor stimulation (in contrast to the majority of microRNAs) (*30*), and because *miR-21* has been implicated in Btg2 regulation in other contexts (*27, 28*). To test whether *miR-21* is responsible for Btg2’s translational repression after acute growth factor stimulation, we generated cell lines that overexpress *miR-21* or an antisense “sponge” to titrate away endogenous *miR-21*, reasoning that these constructs should have opposite effects on YFP accumulation. We created lentiviral expression vectors containing the U6 promoter driving either *miR-21* or the anti-sense sponge, followed by a constitutive CMV promoter driving the expression of TagBFP-NLS to label cells that were successfully transduced. We found that SynIEG cells transduced with the *miR-21* sponge exhibited YFP induction after serum stimulation, whereas mock-transduced or *miR-21*-transduced cells failed to accumulate YFP accumulation (**Figure 3H**). Together, these data confirm a model whereby growth factor signaling drives *BTG2* transcription but *miR-21* blocks Btg2 translation from this mRNA. More broadly, our data suggests that SynIEGs provide a flexible system for studying the decoding of cell signaling stimuli at multiple steps along the central dogma, from transcriptional induction (*e.g*., transcriptional kinetics *via* MS2/MCP imaging) to translational regulation (*e.g*., testing different 5’ and 3’ regulatory sequences) to protein-level regulation (*e.g*., Erk-dependent stabilization of the Fra1 degron).

### A novel AND-gate SynIEG responds to the combination of mitogens and DNA damage

In addition to dissecting the regulation of natural immediate-early genes, SynIEGs are also well-suited for engineering signaling-responsive circuits with novel, desired response functions. To explore this possibility, we set out to develop a SynIEG which implements a signaling-response function that has not been previously described for any immediate-early gene: an AND gate in which growth factor stimulation and a second stimulus, DNA damage, are both co-required to induce gene expression.

We took advantage of *FOS* and *BTG2* regulatory elements whose functions could be composed to produce this AND-gate logic. We previously observed that the *FOS* gene is transcribed in response to growth factor stimulation, regardless of whether DNA damage is present (*14*) (**Figure 4A**). In contrast, 3’ UTR-based translational repression of the *BTG2* gene is abolished in cells exposed to DNA damage either delivered alone or in combination with serum (*14*) (**Figure 4A**). We reasoned that combining these two regulatory elements could result in a circuit that co-requires both serum and DNA damage to promote transcription and relieve translational repression, respectively (**Figure 4B**). We constructed an AND-gate SynIEG with the *FOS* promoter, *FOS* 5’ UTR, *BTG2* 3’ UTR, and a coding sequence driving expression of a destabilized GFP (dGFP) variant that was chosen to achieve a more potent decrease upon the switch to serum-free media and increase the fold-change in GFP induction after stimulation (**Figure 4C**). A constitutive CMV promoter driving TagBFP expression was placed on the same genetic construct to enable fluorescence-based selection of PiggyBAC-transduced NIH3T3 cells. We then performed single-cell sorting to derive two independent clonal cell lines expressing the *fos*-dGFP-*btg2* SynIEG.

We characterized mRNA- and protein-level responses from cells expressing the AND-gate SynIEG in response to three classes of stimuli: doxorubin (DNA damage alone), 1% serum (growth factor alone), or doxorubicin + 1% serum (DNA damage AND growth factor). At the transcriptional level, we observed pronounced MCP foci in *fos-btg2* SynIEG-expressing cells when treated with serum regardless of whether doxorubicin was present (**Figure 4D**), a result that was consistent with our prior observations from the *FOS* endogenous locus (*14*). At the protein level, neither doxorubicin nor serum alone was able to induce dGFP accumulation, whereas their combination triggered dGFP accumulation (**Figure 4E; Figure 5SA**). We further verified that the overall change in dGFP fluorescence (area under the curve; AUC) was significantly increased only by the combination of serum and doxorubicin in both independently-derived clonal cell lines (**Figure 4F; Figure S5B**) and that a control SynIEG (*fos-fos*) did not exhibit AND-gate logic (**Figure S5C**). Taken together, these data demonstrate the successful construction of a stimulus-responsive AND gate using immediate-early gene components. Expression of a desired genetic payload is enabled only when growth factor stimulation induces SynIEG transcription and DNA damage relieves microRNA-mediated translational repression. We emphasize that this AND-gate logic has not been reported for any existing endogenous immediate-early gene, indicating that the SynIEG platform can be used to engineer novel signal-response functions in addition to dissecting endogenous IEG regulation.

## Discussion

There is still a critical gap in our understanding of mammalian signal decoding: how target genes interpret complex combinatorial and dynamic inputs from upstream signaling pathways (*4*). To address this gap, we set out to build mammalian gene cassettes that allow one to simultaneously monitor the transcription and translation of a gene while modulating its various components (i.e. degrons, UTRs, promoter etc.) (**Figure 2A**). After optimizing gene delivery, we quantitatively characterized SynIEG transcription to ensure that their transcriptional regulation was similar to endogenous IEGs (**Figure 2C-D, Figure S2, Movie S1**). We then used SynIEGs to recapitulate the response of the *FOS* gene to dynamic Ras/Erk stimuli (**Figure 3A-D**) and to dissect how *BTG2* responds selectively to DNA damage but not growth factor stimulation despite similar transcriptional behavior in each case (**Figure 3E-H**). In both cases, our experiments highlight regulatory links between signaling pathways and downstream target gene expression that act on multiple nodes of the central dogma (*e.g*., induction of *BTG2* transcription but translational inhibition caused by the *BTG2* 3’UTR), requiring the ability to monitor cellular responses at both transcriptional and post-transcriptional steps. Future studies using SynIEGs can enable the characterization of the signaling pathway to gene expression interface and will shed light on how cells interpret complex biochemical cues to induce a variety of cell fates.

Based on these results, we conjectured that immediate-early genes might also serve as a useful engineering substrate for constructing mammalian cell signaling decoders with desired, novel stimulus-response relationships. Indeed, we found that by combining elements from known immediate-early genes – the *FOS* serum-responsive promoter and the *BTG2* DNA damage-responsive 3’UTR – it was possible to construct a synthetic AND gate that only triggers a 4-fold increase in protein levels in response to combined serum stimulation and DNA damage (**Figure 4**). A striking feature of this AND gate is its simple construction from only two elements, a promoter and 3’ UTR from two separate endogenous IEGs, without additional fine-tuning or optimization. This simplicity follows from the fact that both elements are regulated at distinct steps of the central dogma. SynIEG protein accumulation occurs only if transcription is activated (*via* growth factor stimulation) and translation is de-repressed (*via* DNA damage and micro-RNA regulation). In contrast, engineering AND-gate logic at a single step (e.g. engineering two transcription factors to be mutually required for transcription) often requires complex engineering of three-body interactions (two protein domains with one another and a DNA sequence) (*31*). In addition to constructing logic gates, multi-step regulation is likely to be essential for selectively responding to specific signaling dynamics. For instance, feed-forward loops acting on different timescales have been shown to play crucial roles in discriminating sustained from transient stimuli (*9, 32*).

We propose that SynIEGs implementing novel decoding functions should find utility as reporters of complex endogenous signaling conditions (e.g. marking cells in which some Pathway 1 and Pathway 2 are both active), or as circuits to control the function of engineered cells (*33*). It is known that immediate-early genes are expressed broadly in many tissues, and have extremely potent and fast signaling-induced responses, making them likely to be expressed highly in many contexts (*34, 35*). Furthermore, our work has revealed that SynIEGs display homogeneous transcriptional responses after growth factor stimulation, even after random insertion throughout the genome using the PiggyBAC transposase (**Figure 2D-E, Figure S2**).

This corroborates recent work in which local chromatin structure was seen not to affect transcription of individual genes (*e.g*. actin) (*36*), and where gene expression was largely insensitive to some large-scale genome rearrangements (*37*). All of this suggests that SynIEGs might indeed serve as a predictable platform for mammalian signaling-induced response regulation, but future studies will need to look carefully at other classes of genes before such engineering efforts are applied more broadly. As the field of synthetic biology gets closer to medical applications, especially in the burgeoning field of immunotherapy, cells will need to be engineered to interpret increasingly complex extracellular information (*33*). SynIEGs implementing multi-step regulatory relationships could provide a valuable tool for achieving this goal.

## Methods

### Plasmid Construction

We cloned all of our constructs/synthetic gene circuits into previously published pHR lentiviral expression plasmid (*2*) or into Piggybac plasmid (*38*). For large PCR products (>7500 bp), GXL polymerase was used, followed by overnight DPN1 digestion while for smaller PCR products, HiFi polymerase from Clontech was utilized. Sequences for pFOS, *fos* 5’ UTR, *fos* 3’ UTR, *btg2* 5’ UTR, *btg2* 3’ UTR, *tubulin* 3’ UTR (can be found in supplementary sequences) were obtained via PCR from genomic DNA obtained from NIH 3T3 cells made using Epicentre’s QuickExtract. Sequences for BFP, YFP, MS2 loops and OptoSOS were obtained from plasmids published previously (*14, 39, 40*). Destabilized GFP (dGFP) was obtained as a generous gift from the Reya lab (Addgene: 14715). Fra1 degron was based of FIRE reporter (*20*), which was encoded within an IDT gBlock (sequence can be found in supplementary sequences). PCR products were then run on an agarose gel, purified using Takara Bio’s Nucleospin gel purification kit. Final plasmids were constructed using Takara Bio’s Infusion reagent, amplified in Stellar chemically competent *E. coli* and DNA was extracted using Qiagen miniprep kit.

Construction of a microRNA expression plasmid was made based off of lentiCRISPR v2 plasmid (*41*) in which the U6 promoter is followed by AgeI and EcoRI restriction enzyme cut sites which results in a 7125 bp and 1875 bp band on a agarose gel. The larger piece was excised and purified. miR21 sequence (*agttgtagtcagactattcgat*) and miR21 inhibitor sequence (*tcaacatcagtctgataagcta*) were encoded in duplexed primers with CCGG 5’ overhang for the top strand and AATT 5’ overhang for the bottom strand. T4 ligase was then used to ligate the microRNA sequences into backbone of the plasmid. The resulting plasmid was transformed into Stellar chemically competent *E. coli* and DNA was extracted using Qiagen miniprep kit.

All plasmid verification was done by restriction enzyme digestion and Sanger sequencing by submitting through Genewiz. All plasmids used in this study can be found in Table S1.

### PiggyBac Integration

NIH 3T3s to be integrated were plated 24 hours prior to transfection. 2.08 μg of the plasmid to be integrated along with 0.41 μg of the piggybac helper plasmid were co-transfected into the cells using Liptofectamine LTX with Plus reagent. Cells were selected using FACS for YFP expression after waiting at least 3 days post-transfection. SynIEGs from figures 2 and 3 were integrated into the “Chassis” clonal line described in (*14*). Briefly, the Chassis clonal line is a NIH 3T3 cell line with BFP-Erk, MCP-mCherry, and H2B-iRFP. The *fos*-dGFP-*btg2* synIEG was integrated into wild-type NIH 3T3s. Cells then underwent single cell sorting to isolate clonal cell lines bearing the various synIEG variants.

### Lentivirus production

HEK 293Ts were plated in a 6 well plate at ~40% confluency at least 12 hours before transfection. The cells were then co-transfected with 1.5 ug of the pHR vector of interest along with 1.33 μg and 0.17 μg of CMV and pMD packaging plasmids, respectively using Fugene HD (Promega). Virus was collected after 48 hours post-transfection and filtered through a 0.45mm filter. To the ~2 mL of viral media, 2 μl of polybrene and 40 μl of 1M HEPES were added. Cells to be infected were plated at 40% confluency in a 6 well plate at least 12 hours before infection and then 200-500 ul of viral media was added the cells. 24 hours post transduction virus containing media was replaced with fresh media. Cells were then incubated for at least another 24 hours before sorting using FACS Aria or being placed on the microscope for experiments.

### Cell line maintenance and preparation for imaging

NIH 3T3s were grown in DMEM plus 10% FBS in Thermo Fisher Nunc Cell Culture Tissue Flasks with filter caps at 37C and 5% CO_2_. Cells to be imaged were plated into InVitro Scientific’s 96 well, black-walled, 0.17mm high performance glass bottom plates. 10μg/ml of fibronectin diluted in PBS was placed on the wells, washed off and then cells were plated in DMEM with 10% FBS at least 12 hrs prior to imaging. Between 4 and 6 hours prior to imaging, cells were placed in serum-free media (DMEM with 0.00476mg/mL HEPES). 50μL of mineral oil was pipetted onto the wells right before placing onto the scope to prevent media from evaporating.

### Imaging and optogenetic stimulation hardware

Cells were maintained at 37°C with 5% CO_2_ for the duration of an imaging experiment. Confocal microscopy was performed on a Nikon Eclipse Ti microscope with a Prior linear motorized stage, a Yokogawa CSU-X1 spinning disk, an Agilent laser line module containing 405, 488, 561 and 650 nm lasers, 60x oil emersion objective and an iXon DU897 EMCCD camera.

For optogenetic microscope experiments, blue light from the XLED1 system was delivered through a Polygon400 digital micromirror device (DMD; Mightex Systems) to control the temporal dynamics of light inputs. We applied specific temporal patterns to an image by drawing ROIs within the Nikon Elements software package and using custom macros to turn on and off the light. To attenuate 450 nm light, we dithered the DMD mirrors to apply light 50% of the time, and set our 450 nm LED to 50% of its maximum intensity.

### Drug Treatments

Drug additions were done with a 200 μL gel loading pipette directly onto cells while they were on the microscope. Drugs were pre-diluted to a 1:10 stock concentration additions. Final concentrations for drugs were: cycloheximide (100 μg/ml), doxorubicin (860nM), FBS (10% by volume for all experiments except for the AND-gate experiments which was done at 1% by volume).

### Transcriptional Burst Analysis

Bursting MCP foci were imaged and quantified using a protocol adapted from (*14*). Briefly, 7 z-stack slices spanning 4.5 μm (0.8μm between z-slices) which was centered on the middle of the nucleus. This z-stack was max projected to allow all of the bursts to be visualized on a single plane. Positional information was tracked using the measure tool in Fiji. MATLAB code was used to take in the positional information, fit a 2-dimensional Gaussian to the identified region and finally calculated the integrated area under the fitted Gaussian as the burst intensity. This code can be found in supplementary MATLAB file.

### Microscopy Data Analysis

ND2 files from Nikon Elements software were imported into ImageJ. The measure tool was used to quantify mean intensity of the nuclei of cells of interest. These files were saved and then imported into R to do statistical analysis and graphing. Area-under-the-curve analysis was done by first averaging the first two timepoints to set the baseline. The area was then calculated by using the following formula:

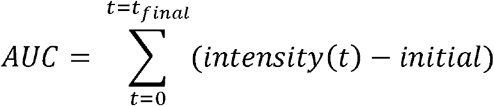

The code for this analysis can be found in Supplementary code.

### Statistical Analysis

Data figure legends include whether data is represented as mean ± S.E.M, mean + S.D or mean ± S.D as well as sample size. For AND gate experiments, the one-sided Student’s t test was applied between serum and the combination of serum and doxorubicin. One-sided student’s t test was deemed appropriate because no prior experiments suggested that doxorubicin would cause decrease in protein accumulation for this particular circuit.

## Author contributions

Conceptualization: P.T.R., M.Z.W. and J.E.T.; Methodology: P.T.R. and J.E.T.; Investigation: P.T.R., and S.G.J. Data analysis: P.T.R., M.Z.W., J.E.T.; Writing – Original Draft: P.T.R. and J.E.T.; Writing – Review and Editing, all authors; Funding Acquisition: J.E.T.; Resources: J.E.T.; Supervision: J.E.T.

## Acknowledgements

We thank all members of the Toettcher lab for helpful comments. We especially thank Dr. John Albeck (UC San Diego) for advice on the Fra-1 degron. We also thank Katherine Rittenbach and Dr. Christina DeCoste of the Princeton Molecular Biology Flow Cytometry Resource Center for cell sorting. This work was supported by NIH grant DP2EB024247 (to J.E.T.), an Innovation Award from the New Jersey Health Foundation (to M.Z.W.), NIH Fellowship F31AR075398 (to S.G.J) and the Lidow Independent Work Research grant and Round Table Fund for Summer Research grant (to P.T.R.).

