## Supplementary Information for "Engineering combinatorial and dynamic decoders using synthetic immediate-early genes"

**This PDF file includes:**

Supplementary Sequences

Supplementary Table 1

Supplementary Figures 1-5

Supplementary Movie 1-2 Legends

**Supplementary Sequences**

*fos* 5’ UTR

cagcgagcaactgagaagactggatagagccggcggttccgcgaacgagcagtgaccgcgctcccacccagctctgctctgcagctcccaccagtgtctacccctggaccccttgccgggctttccccaaacttcgac

*fos* 3’ UTR

gcagtcagagaaggcaaggcagccggcatccagacgtgccactgcccgagctggtgcattacagagaggagaaacacgtcttccctcgaaggttcccgtcgacctagggaggaccttacctgttcgtgaaacacaccaggctgtgggcctcaaggacttgcaagcatccacatctggcctccagtcctcacctcttccagagatgtagcaaaaacaaaacaaaacaaaacaaaaaaccgcatggagtgtgttgttcctagtgacacctgagagctggtagttagtagagcatgtgagtcaaggcctggtctgtgtctcttttctctttctccttagttttctcatagcactaactaatctgttgggttcattattggaattaacctggtgctggattgtatctagtgcagctgattttaacaatacctactgtgttcctggcaatagcgtgttccaattagaaacgaccaatattaaactaagaaaagataggactttattttccagtagatagaaatcaatagctatatccatgtactgtagtccttcagcgtcaatgttcattgtcatgttactgatcatgcattgtcgaggtggtctgaatgttctgacattaacagttttccatgaaaacgtttttattgtgttttcaatttatttattaagatggattctcagatatttatatttttattttatttttttctaccctgaggtctttcgacatgtggaaagtgaatttgaatgaaaaattttaagcattgtttgcttattgttccaagacattgtcaataaaagcatttaagttgaa

*btg2* 5’ UTR

gagtggtatgaaaggcgcagcccggggaaagtccgggcagagcccgagaggtggccagaccgtcatcatcgttctaatacagctacttcctcagccaccggt

*btg2* 3’ UTR

ataggagccacccgaccctggcactctactgttctcatgctgccctgacaacaggccaccgtatacctcaacctggggaactgtatttttaaaatgaagagctatttatacatgttttttttttttttttttaagaaaagaggaaaaaaaccaaaagattttttttaaaaaaaaaagaaaaagaaaaacaattctttaaagggagctgcttggaagtggcctccccaggtgcctttggagagaactgttcttgattgcatctgtgagccagtgtttgcctaggggaatgggttggggattggcctagccaaggtaaaaggggattcttggctgatccccccaggaggtggtggaagggagcaaggttagcaactgtgaatgagaggggtcagggtctgctctgggttaccgtcccagctgggatgcctgtatgcctggtccctctcttactcaggggcattcaagcctgatcttaaataatactacattgcctaatcttctcttttgtttttcggctgagatcaggggcagactgaaaggcctctcctgtcccttctgttctaagcagtctcttgaagccgtgtctcgtttctgagtctacccttgggggcctgaagagcttgcttcccagcccggaatctgtctaaacattttttggagggtgggatgtaaggcaggcgcgcaaagactttggggctaagatggagaggacctgcacaaaacctttgctttgctctgtgctgctttgtatgggtggatagtgaataatttagggatgatttgcaatggaattttgggacccaaagagtatccaatgggggtaggtcttttggacccagtccttccttttgggaaccacacgacagtctgaatgctgctaccactattcctttgagaggtggctcaaagctccagggaactccaagtcctttcttactgccttctcttcaagagcaaccttccccattttttctttcccctttcctgcggttgggtcctggagggccctatttcctaggacaagcgttctcagtcactgtgcaatagtcccaggatgctctgagaccggacctcccagcccctcctgatgccctggtaggttttagggacccattcttcccattcttctaggggttctgactggctggtgggccttgcggagatcttcctgggccacagggagggcacctgtgcactgcaggactacctggtattcttgtagggctgccatgaagagtcaaaccttgggcacagctttagctccttggtgctcagagcacctgtgggggaggttacctctctctctctcttagtaaaatccaaatttatttgtagatgtgtgcaatatttactgttctgggttggagaaaattgggaaacactgggaagaaatggctttccttcaggttcagtgacactgatgagggcttcttagaaggcctcaagtctctcaaactgaaggacagagctagagctagcccgtcaccccttggtgaggattcccttccccgtttctctccaccgccatggcatcctgtgtcctagatttctcagctcctcagtttctgctcaaaggtgctatttaccaaactctctgcctgcccgggcagacaggccccagcttcgcacagccttcccaggtggcttcgtctctcttgctttaaccttaactctgggcccacagacctgagagctgtggcctacaacaaagctgtgaattgttccagatggttcttgtgttttgtccgcacacaggtgcctgccgtttagaagctgcctcctggtctcatgcttaaatcttcaattctttactgtccttgttcactttagaaatgacaaaccctagagctggactgttgagcaggcctgtctctcttattaagtaaaaataagaaatagtggtaagtttgtaagctattctgacagaaaagacaaaggttactaattgtataatagcgcttttatatggaagactgtacagctttatggacaaatgtaaactttcttttttgtttttaataaaaatgtagcagatcgtgtaatgtgtagagaaggtggaattccatagcgctgactggccctcttagatacaaaccccttcgtgtttcgtggacttcatcagttgtcctgagctccgtctgcagcatgcagacgttataaaagataccctaggtttgtgacgaggttgcttccattgcatcccctctccttgggaggcggctccatacttctgtccttgtgaatattgtcacaggtctcttaaaaaacaaacaaacaaaaaagcccaaaaaccaatatgttatctgcacactctgcagccagttccatagttttgctcttggccattcagcacattgaagagctggccagctgtgtccacatctgcgcaagcaagccaccacggctggaaactatataaagaaaagaaaccatccagaaagttccaaagacacaagacttgggtttggcacacactaagcacaccagtgcactcgatgcgactg

*tubulin* 3’ UTR

ttcttaaagcttttactttgagacatcatggaaaacttaagaggtacaacatggagaagacatgatcacagaatggaaacagcacagaagcatcagtgcctgcaactaatactggagcagtttgacgacacagggggctcaaggaatggacttagtactcctctccttcttcctccccctcgccttccacagaatccacaccaacctcctcataatccttctctagggcagccatgtcctcacgggcctcagagaactcaccctcctccatgccctcacccacataccagtgcacaaaggcacgcttggcatacatcagatcaaacttgtgatctaggcgagcccaggcctcagcaatggctgtggtgttgctcagcatgcacacagctctctgcaccttggccaggtcaccaccgggtaccacagtgggaggctggtaattaatgccaaccttgaagccagtggggcaccagtctacaaactggatgctgcgcttggtcttgatggtggcaatggcagcattgacatctttgggaaccacatcaccacggtatagcaggcagcaagccatgtatttaccatggcgagggtcacatttcaccatctggttggctggctcaaagcaggcattggtgatctctgctacagaaagctgctcatggtaggctttctcagcagagatgacaggggcataagtggccagagggaagtggatgcgagggtagggtaccaggttggtctggaattctgtcagatcaacattcagggccccatcaaatctgagggaagcagtgatggaagacacaatctggctaataaggcggttaaggttggtgtaggttgggcgctcaatgtcgaggtttctacgacagatgtcatagatggcctcattgtctaccatgaaggcacaatcagagtgctccagggtggtgtgggtggtgaggatggaattgtagggctcaaccacagcagtggaaacctggggggctgggtaaatggagaactccagcttggacttctttccgtaatccacagagagccgctccatcagcagggaggtgaagccagagccagttcccccaccaaagctgtggaaaaccaagaagccctggagacctgtgcactggtcagc

Fra1 degron amino acid sequence

LVLEAHRPICKIPEGDKKDPGGSGSTSGASSPPAPGRPVPCISLSPGPVLEPEALHTPTLMTTPSLTPFTPSLVFTYPSTPEPCSSAHRKSSSSSGDPSSDPLGSPTLLAL

**Table S1: Resources and Plasmids**

| REAGENT or RESOURCE | SOURCE | | | IDENTIFIER |
| --- | --- | --- | --- | --- |
| Bacterial and Virus Strains |  | | |  |
| Stellar Chemically Competent Cells | Clontech Laboratories | | | Cat# 636763 |
| Chemicals, Peptides, and Recombinant Proteins | | |  |  |
| CloneAmp HiFi PCR Polymerase | Clontech Laboratories | | | Cat#639298 |
| Cycloheximide | Sigma-Aldrich | | | Cat# C104450 |
| Doxorubicin | Fischer Scientific | | | Cat# ICN15910101 |
| Trichostatin A | Tocris | | | Cat# 1406 |
| Fugene HD | Promega | | | Cat# E2311 |
| Lipofectamine LTX with Plus Reagent | Thermo-Fischer | | | Cat# 15338100 |
| PrimeSTAR GXL DNA Polymerase | Clontech Laboratories | | | Cat# R050B |
| QuickExtract DNA Extraction | Epicenter Bio | | | Cat# QE09050 |
| Experimental Models: Cell Lines | |  | |  |
| NIH 3T3 Cells | ATCC | | | Cat# CRL-1658 |
| Lenti-X 293t Cells | Clontech Laboratoires | | | Cat# 632180 |
| ‘‘Chassis’’ cell line (NIH 3T3 + optoSOS + H2B-diRFP + MCP-mCherry ) | Wilson et. al., 2017^16^ | | | Available upon request |
| Recombinant DNA |  | | |  |
| pCMV-dR8.91 lenti helper plasmid | Gift from the Trono lab | | | Available upon request |
| pMD2.G lenti helper plasmid | Gift from the Trono lab | | | Addgene #12259 |
| pHR *fos-tubulin* synIEG | This paper | | | Available upon request |
| PigBac *fos-tubulin* synIEG | This paper | | | Available upon request |
| PigBac *fos-fos* synIEG | This paper | | | Available upon request |
| PigBac *fos-btg2* synIEG | This paper | | | Available upon request |
| PigBac *btg2-fos* synIEG | This paper | | | Available upon request |
| PigBac *btg2-btg2* synIEG CMV BFP | This paper | | | Available upon request |
| PigBac *fos*-dGFP-*btg2* synIEG CMV BFP | This paper | | | Available upon request |
| PigBac *fos*-YFP-Fra1-*fos* synIEG | This paper | | | Available upon request |
| pHR U6 “*empty”* CMV BFP-NLS | This paper | | | Available upon request |
| pHR U6 miR-21 CMV BFP-NLS | This paper | | | Available upon request |
| pHR U6 miR-21 inhibitor CMV BFP-NLS | This paper | | | Available upon request |
| pHR BFP-SSPB-SOScat p2a iLID-CAAX | This paper | | | Available upon request |
| pLenti TOPFLASH d2EGFP | Gift from Reya lab | | | Addgene #14715 |

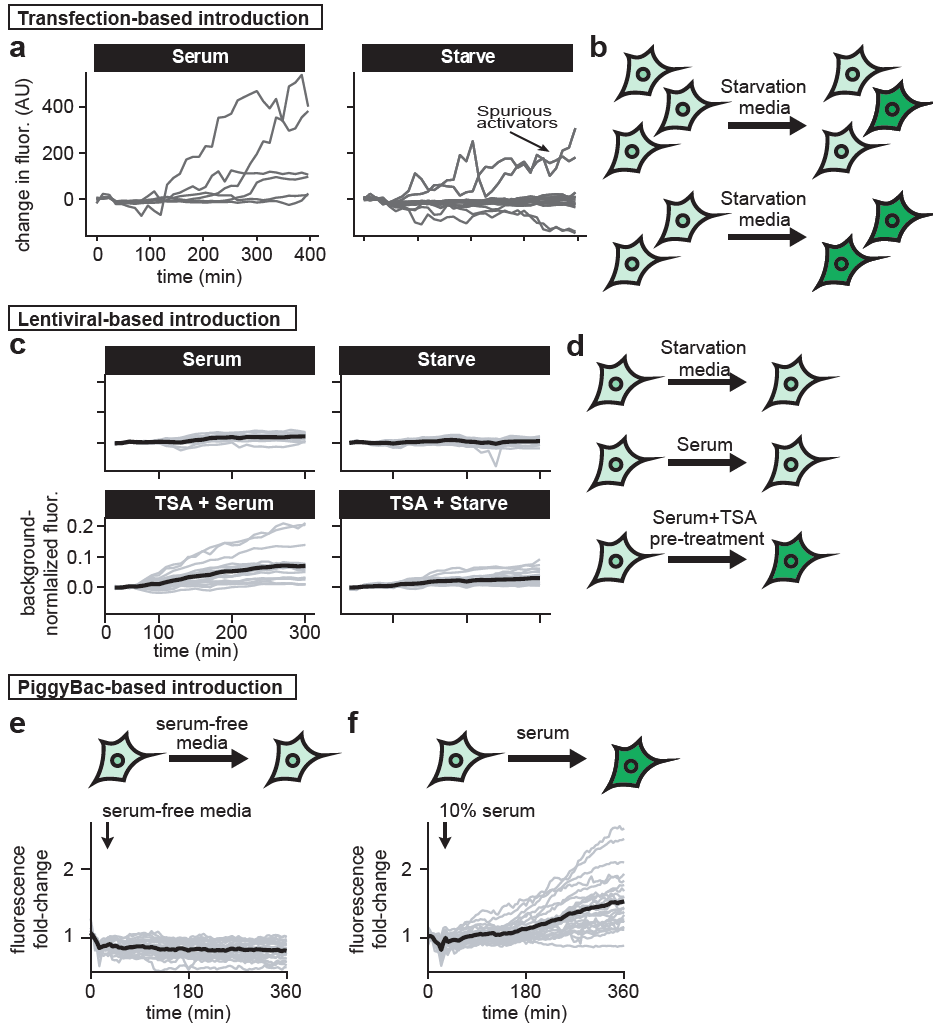

**Figure S1. Comparison of methods to introduce synthetic target genes into target cells.** (**A**) Quantification of YFP fluorescence in NIH 3T3 after being transfected with *fos-tub* synIEG and treated with either serum (n=6 cells) or growth factor free media (n=20 cells). Arrow denotes some of the traces in the starvation case in which YFP intensity increases. (**B**) Model of transfection based method of delivery for synIEGs. Also serum induced cells increase in YFP fluorescence, there is spurious activation of synIEGs in the serum free case. (**C**) Dynamic traces of NIH 3T3s that were lentivirally integrated with *fos-tubulin* synIEG. Means (dark black line) and single cell traces (light gray lines) are shown for cells that were either given starvation media (n=16 cells), serum stimulation (n=10 cells), pre-treated with TSA and then given starvation media (n=20 cells) or pre-treated with TSA and then treated with serum (n=19 cells). (**D**) Model for lentiviral transduction of synIEGs: after silencing occurs, neither starvation media nor serum induces YFP accumulation but with the addition of trichostatin A, YFP is now inducible. (**E-F**) Dynamic single cell traces (light gray lines) with mean shown in dark black line for NIH 3T3s containing the *fos-tubulin* synIEG after PiggyBac integration for cells that received starvation media (in **E**; n=27 cells) and serum (in **F**; n=26 cells).

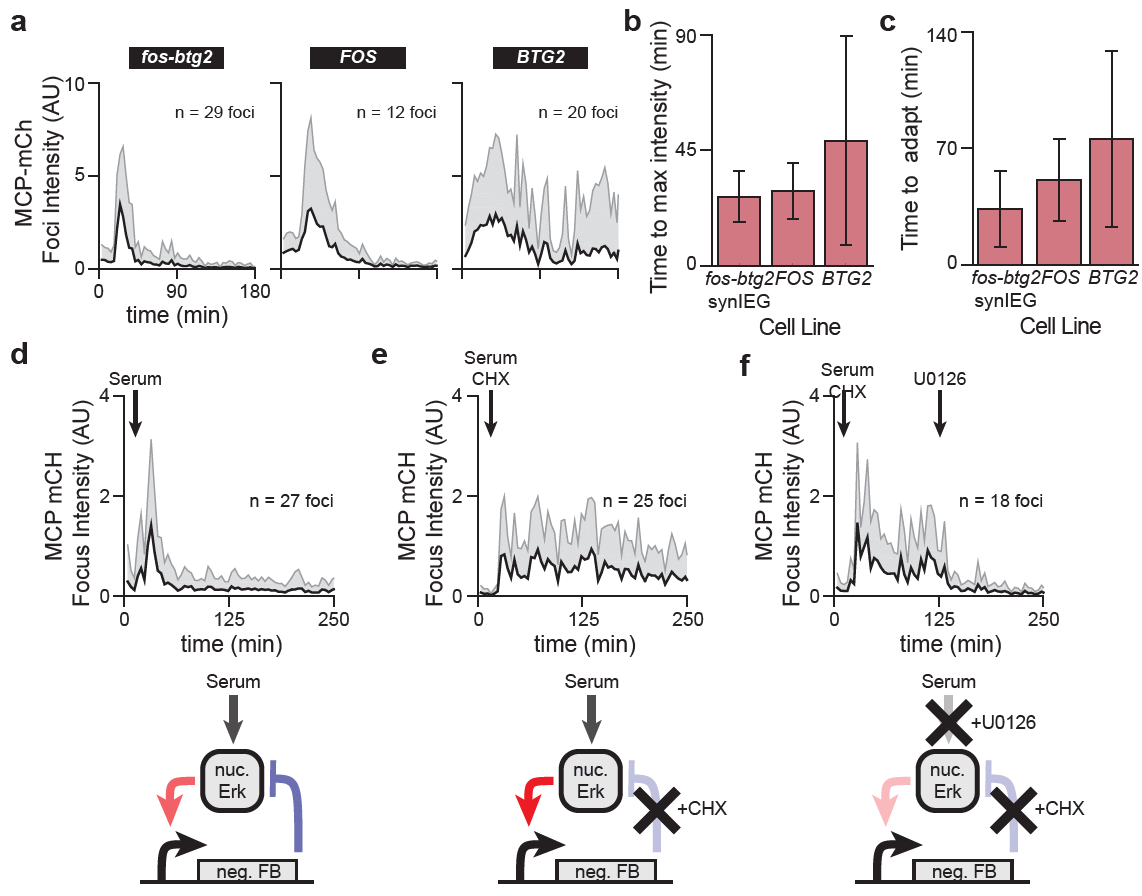

**Figure S2. synIEGs recapitulate the dynamics and regulation of endogenous IEGs.** (**A**) Mean + S.D for transcriptional foci intensity after serum stimulation for NIH 3T3s either containing YFP-MS2 tag on endogenous *BTG2* (n = 20 foci) or *FOS* (n= 12 foci) endogenous gene or *fos-btg2* synIEG (n = 29 foci). (**B**) Bar graph showing mean± S.D for the time until maximum intensity of transcriptional foci after serum stimulation for cells quantified in **A**. (**C**) Bar graph showing mean± S.D for the time until adaptation (defined as the time after reaching maximum intensity when the burst intensity is 50% of the maximum) for cells quantified in **A**. (**D-E**) Quantification of MCP-mCherry nuclear foci (mean + S.D.) in fos-btg2 synIEG cells treated with serum (in **D**; n= 27 foci), serum + cycloheximide (in **E**; n=25 foci) or serum and cycloheximide followed by addition of the MEK inhibitor U0126 after 90 min (in **F**; n= 18 foci). Underneath each graph is a schematic of the interactions occurring between nuclear Erk, and production of its negative feedback.

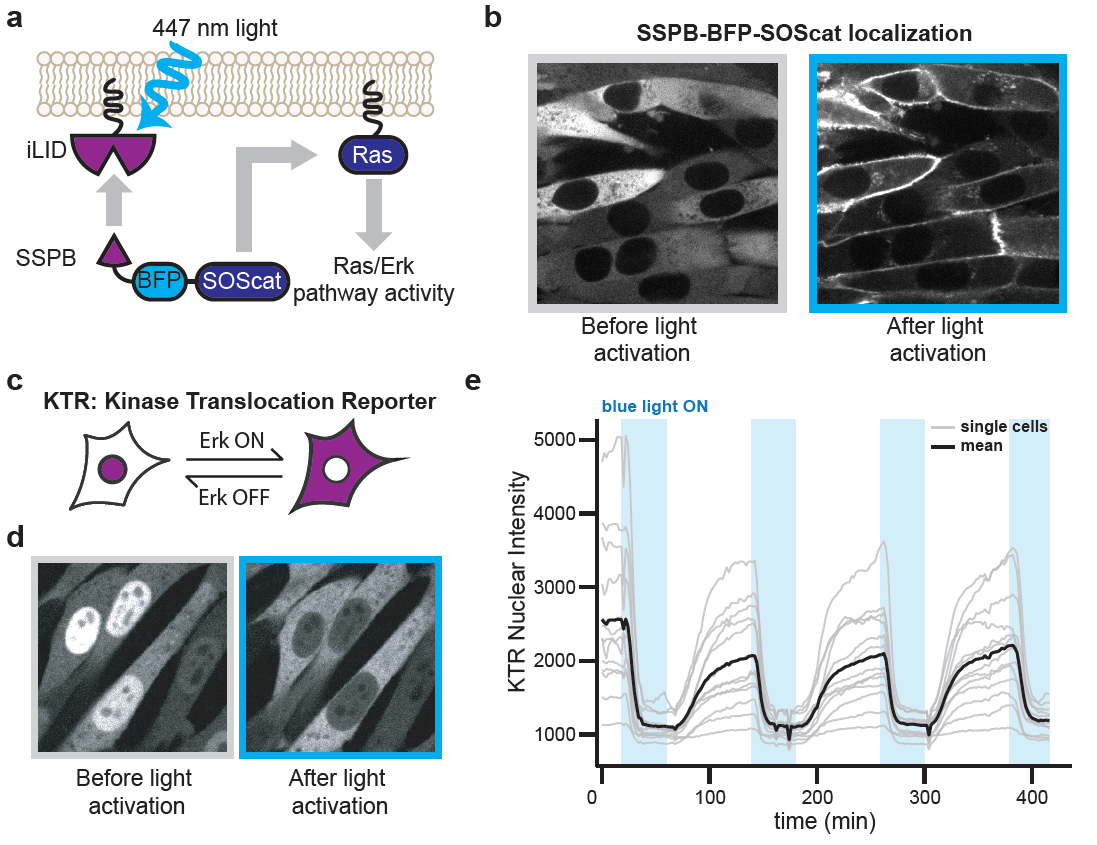

**Figure S3. The iLID-OptoSOS system allows for transient and sustained activation of MAPK activity in SynIEG-expressing cell lines.** (**A**) Diagram of the OptoSOS system where blue light dimerizes iLID and SSPB, recruiting SOScat to the membrane, activating the Ras-Erk pathway. (**B**) Representative image of BFP-SOScat before and after blue light stimulation. (**C**) Schematic of the kinase translocation reporter (KTR) that is present in the nucleus when Erk is off and in the cytoplasm after Erk is on. (**D**) Representative images of the iRFP channel for NIH 3T3s expressing the OptoSOS – iLID/SSPB system as well as KTR fused to iRFP. Before light, cells have Erk off as evidenced by nuclear KTR and after blue light stimulation, cells have turned Erk on. (**E**) Dynamic traces of 13 NIH 3T3 cells expressing the OptoSOS – iLID/SSPB system as well as KTR fused to iRFP where dark black line represents the mean and gray traces are individual cells. Blue bars show where the blue DMD was turned on. Cells were given 10 minutes no light, 20 minutes blue light and then 30 minute no light cycles over the course of the experiment.

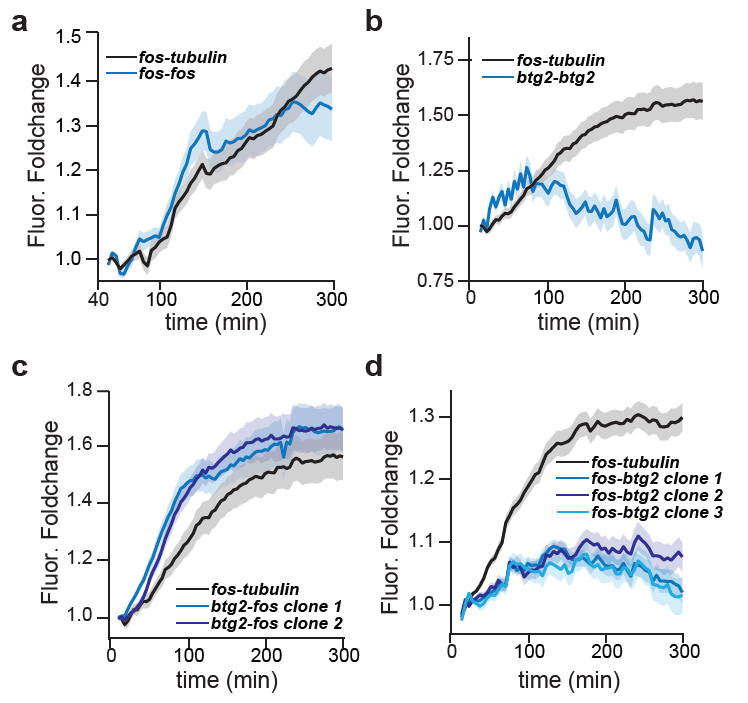

**Figure S4. The *BTG2* 3’ UTR is necessary and sufficient for translation inhibition.**
(**A-D**) Quantification of YFP induction from clonal NIH3T3 cell lines expression each of four SynIEGs: *btg2-btg2* (in **A**), *fos-btg2* (in **B**), *btg2-fos* (in **C**), or *fos-fos* (in **D**). In each case, YFP induction was compared to a control *fos-tubulin* SynIEG. Means ± S.E.M are shown for at least 20 cells in each condition.

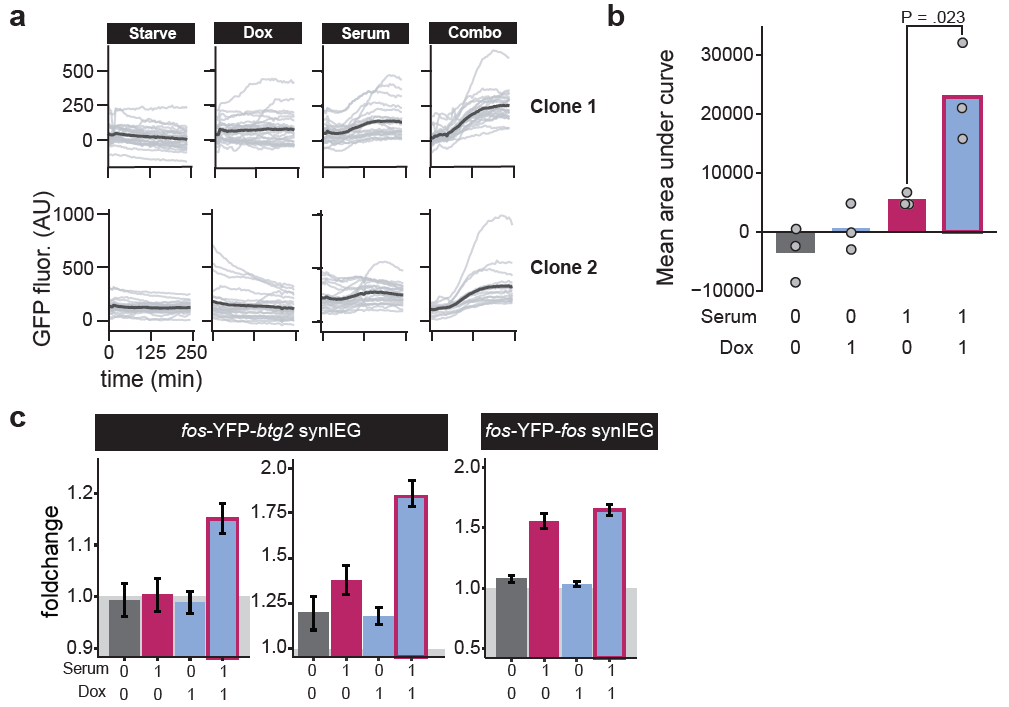

**Figure S5. Further characterization of the *fos-btg2* AND gate.** (**A**) GFP induction traces for clonal NIH3T3 cells harboring the *fos-*dGFP*-btg2* synIEG. Cells were treated with growth factor free media, 860 nM doxorubicin (Dox), 1% serum, or a combination of doxorubicin and serum. Gray traces are single cells and dark black line is the mean. (**B**) Quantification of ~20 cells for each condition in 3 biological replicates for clone 2 (see Methods). Each point is the mean of at least 20 cells in a single experimental replicate; the bar represents the mean of three repeats. Statistical analysis was done using the Student’s t-test between serum and combination conditions. (**C**) Fold-change in YFP induction for clonal NIH3T3 cell lines harboring either the *fos-*YFP-*btg2* synIEG (2 independent clones; left panels) or the *fos-*YFP-*fos* synIEG (right panel). Cells were treated with starvation media, 1% serum, 860 nM doxorubicin, or a combination of doxorubicin and serum. Bar height represents the mean YFP fold-change over initial time-point after 300 minutes of stimulation and error bars represent S.E.M for at least 20 cells per condition.

**Supplementary Movie Legends**

**Movie S1.** Time-lapse imaging of NIH 3T3 containing *fos-btg2* SynIEG in the MCP-mCherry channel to visualize transcriptional induction. MCP-mCherry images were collected at 0.7 μm z-stacks and the processed using maximum intensity projection. Cells were stimulated with serum and images were acquired every 3 minutes.
