## Supplementary Code 1 for "Engineering combinatorial and dynamic decoders using synthetic immediate-early genes": analysisScript.m

```
%% SOME PARAMETERS
p.umPerPixel = 4.2221; % Never change this. Scale of pixels at 60X

% Change this to change the intensities on the plots for burst finding
p.PLOT_SCALE = [20 80]*10;

% This is if you are concerned about the size of the region you are
% analyzing. You can "expand" using delta_region (in pixels)
% delta_region = 5; % pixels to grow in each dimension
p.delta_region = 1; % pixels to grow in each dimension


% This is whether you want to plot
p.toPlot = 0; % Do you want to plot??

%% Analyze for Fig 2e
p.nT = 33; % The # of timepoints in your imaging expt. Change as needed!
p.hmin = 700; % h-minima transform to smoothen out noisy crap. 0 = don't use.

% You can specify burst files this way, or in a cell array, or however you want!
pathname = '...Code Deposit\Dataset';
image_file = 'V_BandPass001-MaxIP.nd2 - Z=5 C=0.tif';

burst_files = dir([pathname, '/*.txt']);
burst_files = {burst_files(:).name}';
% burst_files = {'Sq_1-XY_5-C_R-cell_1-spot_5.txt'};
% You can get an image file however you want too

BURST = analyze_bursts(pathname, image_file, burst_files, p);
plot(BURST)
% save fig2e BURST p burst_files image_file;
```
