## Supplementary Code 1 for "Engineering combinatorial and dynamic decoders using synthetic immediate-early genes": analyze_bursts.m

```
function BURST = analyze_bursts(pathname, image_file, burst_files, p)
%% LOAD THE IMAGES
for i = 1:p.nT
    IM_RFP(:,:,i) = imread([pathname '/' image_file], i);
end

%% MAIN LOOP over our cells
%  Put the inside of this code into its own function

for i = 1:length(burst_files) 
    % Load your hand-drawn circles
    fname = burst_files{i};
    tmp = importdata([pathname '/' fname]);
    
    i_slice = find(~cellfun(@isempty, strfind(tmp.textdata, 'Slice')));
    i_bx    = find(~cellfun(@isempty, strfind(tmp.textdata, 'BX')));
    i_by    = find(~cellfun(@isempty, strfind(tmp.textdata, 'BY')));
    i_W     = find(~cellfun(@isempty, strfind(tmp.textdata, 'W')));
    i_H     = find(~cellfun(@isempty, strfind(tmp.textdata, 'H')));
    slice   = tmp.data(:,i_slice);
    
    for t = 1:length(slice)
        box(slice(t)).bx = round(p.umPerPixel*tmp.data(t,i_bx)) - p.delta_region;
        box(slice(t)).by = round(p.umPerPixel*tmp.data(t,i_by)) - p.delta_region;
        box(slice(t)).W  = round(p.umPerPixel*tmp.data(t,i_W)) + 2*p.delta_region;
        box(slice(t)).H  = round(p.umPerPixel*tmp.data(t,i_H)) + 2*p.delta_region;
    end
    
    for t = 1:length(slice)
        xx  = box(slice(t)).bx+(1:box(slice(t)).W);
        yy  = box(slice(t)).by+(1:box(slice(t)).H);
        Z   = double(IM_RFP(yy,xx,t));
        Z   = imhmin(Z, p.hmin);
        
        %Zdelta = Z(p.delta_region:end - p.delta_region, p.delta_region:end - p.delta_region);
        %zvd    = Zdelta(:);
        [m n] = size(Z);
        BW = conv2(ones(m-1,n-1), ones(2,2)) == 4; % a ring of zeros
        BW = imerode(BW, strel(ones(5,5)));
        Zdelta = Z.*BW;
        zvd = Zdelta(:);
        
        [X, Y] = meshgrid(xx, yy);

        % Data to fit
        xv = X(:);
        yv = Y(:);
        zv = Z(:);
        
        % Get the initial guesses, maxes & mins for the fit
        [ii, jj] = find(Z == max(zvd), 1, 'first');
        
        a0 = max(zv)-mean(zv); am = 0; aM = 5*mean(zv);
        b0 = mean(zv); bm = mean(zv)-std(zv); bM = mean(zv)+std(zv);
        c0 = 1; cm = 0.4; cM = 1;
        x0 = X(ii,jj); xm = max(x0-3, xv(1)+2); xM = min(x0+3, xv(end)-2);
        y0 = Y(ii,jj); ym = max(y0-3, yv(1)+2); yM = min(y0+3, yv(end)-2);

        fo = fitoptions('Method','NonlinearLeastSquares',...
                   'Lower',[am bm cm xm ym],...
                   'Upper',[aM bM cM xM yM],...
                   'StartPoint',[a0 b0 c0 x0 y0]);
        
        ft = fittype('a*exp(-((x-x0)^2+(y-y0)^2)/(2*c^2)) + b', 'independent', {'x' 'y'}, 'dependent', 'z');
        
        if p.toPlot
            cfit = fit([xv yv], zv, ft, fo)
        else
            cfit = fit([xv yv], zv, ft, fo);
        end
        
        BURST(t,i) = 2*pi*cfit.a*cfit.c^2;
        
        if p.toPlot
            % Plot
            figure(1)
            % set(gcf, 'position', [325 616 1500 315])
            subplot(1,3,1)
            imshow(Z, p.PLOT_SCALE)
            subplot(1,3,2)
            surf(X,Y,Z)
            set(gca, 'zlim', p.PLOT_SCALE)
            subplot(1,3,3)
            zvm = cfit(xv, yv);
            ZM = reshape(zvm, size(Z));
            surf(X, Y, ZM)
            set(gca, 'zlim', p.PLOT_SCALE)
            title(sprintf('burst(i) = %g', BURST(t,i)));
            drawnow
            pause
        end
    end
    if p.toPlot
        % Plot the full trajectory
        figure(2)
        % set(gcf, 'position', [146 784 513 100])
        plot(BURST(:,i))
        pause
    end
    fprintf('.')
end
fprintf('\n')
```
