## Supplementary Code 1 for "Engineering combinatorial and dynamic decoders using synthetic immediate-early genes": generateFigs.m

```
%% Generate Figure 2E
load ('fig2e.mat')
T = (1:33) * 3;
plot (T, BURST,'r')
% sd = std(BURST');
% shadedErrorBar(T, mean(BURST,2), sd(:))
title('Single Cell Bursts')

%% Generate Figure S2C-E
load('figS2Cserumall.mat')
load('figS2Dchxall.mat')
load('figS2EU0126.mat')

T = (1:63)*4;

figure (2)

subplot (1,3,1)
hold on
plot (T, SerumBursts,'Color', [.5 .5 .5]) 
plot (T, mean(SerumBursts,2), 'r', 'LineWidth', 2)
ylim([0 70000])
title('Serum alone')
hold off

subplot (1,3,2)
hold on
plot (T, CHXBursts,'Color', [.5 .5 .5])  
plot (T, mean(CHXBursts,2), 'r', 'LineWidth', 2)
ylim([0 70000])
title('Serum+CHX')
hold off 

subplot (1,3,3)
hold on
plot (T,U0126Bursts,'Color', [.5 .5 .5])
plot (T, mean(U0126Bursts,2), 'r', 'LineWidth', 2)
ylim([0 70000])
title('Serum+CHX+U0126')
hold off 
%% Make FigS2A
load ('figS2ABtg2All.mat')
load ('figS2AV1all.mat')
load ('figS2AFosAll.mat')

T = (1:52)*3.5;

figure (3)
subplot (1,3,1)
hold on
plot (T, Btg2Bursts, 'Color', [.5 .5 .5])
plot (T, mean(Btg2Bursts, 2), 'r', 'LineWidth', 2)
ylim ([0 18000])
title('Btg2')
hold off

subplot (1,3,2)
hold on
plot (T, V1Allbursts, 'Color', [.5 .5 .5])
plot (T, mean(V1Allbursts,2), 'r', 'LineWidth', 2)
ylim ([0 18000])
title('fos-btg2 synIEG')
hold off

subplot (1,3,3)
hold on
plot (T, FosAll, 'Color', [.5 .5 .5])
plot (T, mean(FosAll,2), 'r', 'LineWidth', 2)
ylim ([0 18000])
title('Fos')
hold off

%% Generate FigS2B-C
[m.Btg2,idx]=max(Btg2Bursts);
[m.V1,idx2] = max(V1Allbursts);
[m.Fos,idx3] = max(FosAll);

I = [3.5*idx 3.5*idx2 3.5*idx3];
AverageIdx = [3.5*mean(idx) 3.5*mean(idx2) 3.5*mean(idx3)];
stddevIdx = [3.5*std(idx) 3.5*std(idx2) 3.5*std(idx3)];

thresh = 1;
threshold.Btg2 = thresh * (m.Btg2 + (Btg2Bursts(52,:)))/2;
threshold.Fos = thresh * (m.Fos + (FosAll(52,:)))/2;
threshold.V1 = thresh * (m.V1+(V1Allbursts(52,:)))/2;

for (i=1:size(Btg2Bursts,2))
    temp = Btg2Bursts(:,i);
    timepoint = find(temp > threshold.Btg2(1,i) , 1, 'last');
    if isempty(timepoint)
        timepoint = size(Btg2Bursts,1);
    end
    adaptation.Btg2(i) = timepoint;
end

for (i=1:size(FosAll,2))
    temp = FosAll(:,i);
    timepoint = find(temp > threshold.Fos(1,i) , 1, 'last');
    adaptation.Fos(i) = timepoint;
end

for (i=1:size(V1Allbursts,2))
    temp = V1Allbursts(:,i);
    timepoint = find(temp > threshold.V1(1,i) , 1, 'last');
    adaptation.V1(i) = timepoint;
end

AverageAdapt = [3.5*mean(adaptation.Btg2) 3.5*mean(adaptation.V1) 3.5*mean(adaptation.Fos)];
stddev = [3.5*std(adaptation.Btg2) 3.5*std(adaptation.V1) 3.5*std(adaptation.Fos)];


figure (4)
subplot (2,1,1)
hold on 
bar (AverageIdx)
errorbar(AverageIdx,stddevIdx,'.')
hold off
title('Time to Max')

subplot (2,1,2)
hold on
bar(AverageAdapt)
errorbar(AverageAdapt, stddev,'.')
hold off
title('Time to Adapt')


%% Generate FigS5A
load ('figS5ASerum.mat')
load ('figS5ACombo.mat')
load ('figS5ADox.mat')

T = (1:60) *3;

figure(5)
subplot(3,1,1)
hold on
plot (T, SerumS5bursts, 'Color', [.5 .5 .5])
plot (T, mean(SerumS5bursts,2), 'r', 'LineWidth', 2)
ylim([0 150000])
title('Serum alone')
hold off

subplot(3,1,2)
hold on
plot(T, ComboBursts, 'Color', [.5 .5 .5])
plot (T, mean(ComboBursts, 2), 'r', 'LineWidth' ,2)
ylim([0 150000])
title('Serum + Dox')
hold off

subplot(3,1,3)
hold on
plot (T, DoxBursts, 'Color', [.5 .5 .5])
plot (T, mean(DoxBursts, 2), 'r', 'LineWidth', 2)
ylim([0 150000])
title('Dox alone')
hold off
```
