## Supplementary Code 1 for "Engineering combinatorial and dynamic decoders using synthetic immediate-early genes": bfilt2n.m

```
function y=bfilt2n(x,dim)
% Blaise Filter
if nargin < 2 || dim == 1
    iamvert = 0;
elseif dim == 2
    x = x';
    iamvert = 1;
else
    error('Invalid second argument: DIM must be equal to 1 or 2');
end

xx=x;
clear x;
for i=1:size(xx,1)
    x=xx(i,:);    
    
    l=length(x);
    
    m(1,:)=[x(3) x(2) x(1:l-2)];
    m(2,:)=[x(2) x(1:l-1)];
    m(3,:)=x;
    m(4,:)=[x(2:l) x(l-1)];
    m(5,:)=[x(3:l) x(l-1) x(l-2)];
    
    b=[0 .25 .5 .25 0];
    
    for j = 1:size(xx,2)
        if sum(m(:,j) == 0) > 1
            y(i,j) = xx(i,j);
        else
            y(i,j) = b*sort(m(:,j));
        end
    end
end
if iamvert
    y = y';
end;
```
