## Supplementary Code 1 for "Engineering combinatorial and dynamic decoders using synthetic immediate-early genes": curateNuclei2.m

```
function BWout = curateNuclei(BWin)
warning off

% keyboard
%% Filter out smaller-than-10 objects
BWout = imopen(BWin, strel('disk', 10));

%% Watershed to get single cells
WATER_THR = 0.5;
D = -bwdist(~BWout);
D = imhmin(D, WATER_THR);
L = watershed(D);

% This next step "widens" the lines in the watershed to REALLY separate
% objects
BWout = BWout & imerode(L, strel('disk', 5));

% BWout = imclearborder(BWout);

warning on
```
