## Supplementary Code 1 for "Engineering combinatorial and dynamic decoders using synthetic immediate-early genes": curateNuclei.m

```
function BWout = curateNuclei(BWin)

warning off

R_min = 10; % min radius of 10 pixels per nucleus
A_min = pi*R_min.^2;

%% Now get rid of objects that don't meet the threshold

CC = bwconncomp(BWin);
numPixels = cellfun(@numel,CC.PixelIdxList);
idx = find(numPixels > A_min);

BWout = false(size(BWin));
for i = 1:length(idx)
    BWout(CC.PixelIdxList{idx(i)}) = 1;
end

%% Fill holes & smoothen edges
BWout = imfill(BWout>0, 'holes');
BWout = imclose(BWout>0, strel('disk', 5));
BWout = imopen(BWout>0, strel('disk', 8));

%% Watershed to get single cells
WATER_THR = 0.5;

D = -bwdist(~BWout);
D = imhmin(D, WATER_THR);
L = watershed(D);
% This next step "widens" the lines in the watershed to REALLY separate
% objects
BWout = BWout & imerode(L, strel('disk', 7));

%% Get region props
rp = regionprops(BWout, 'eccentricity', 'area', 'centroid', 'solidity');
[L n] = bwlabel(BWout);
rps = [rp.Solidity];
rpa = [rp.Area];

%% Filter region props
% now also make sure you're approximately round
idx = find(rps > 0.7 & rpa > pi*R_min^2);
notidx = setdiff(1:n, idx);
for j = 1:length(notidx)
    L(L == notidx(j)) = 0;
end

%% BWout is final mask of nuclei
BWout = L > 0;
BWout = imclearborder(BWout);

warning on
```
