## Supplementary Code 1 for "Engineering combinatorial and dynamic decoders using synthetic immediate-early genes": getBackgroundIntensity.m

```
function bkgd = getBackgroundIntensity(IM)
% Subsample image for kmeans
A_IM = size(IM,1)*size(IM,2);
nSamples = min(A_IM, 1e4);
i_samp = ceil(A_IM*rand(nSamples,1));

for i = 1:size(IM,3)
    tmp = IM(:,:,i);
    ivals(:,i) = double(tmp(i_samp));
end

[idx c] = kmeans(ivals, 3, 'emptyaction', 'singleton', 'start', [20;50;100]);
c = sort(c);
bkgd = c(1);

[y x] = hist(ivals,200);
plot(x,y/sum(y))
line(bkgd*[1 1], [0 0.2], 'color', 'k', 'linewidth', 2)
```
