## Supplementary Code 1 for "Engineering combinatorial and dynamic decoders using synthetic immediate-early genes": getCellBoxes.m

```
function IM_cell = getCellBoxes(r_track, c_track, IM)

% BOX SIZE
H = 50;

% IMAGE SIZE (assumes square)
L = size(IM,1);

nT     = size(r_track, 1);
nCells = size(r_track, 2);

%% Get all boxes

for i = 1:nCells
    for j = 1:nT
        if r_track(j,i)-H < 1
            box{i}(j,1:2) = [1 (2*H+1)];
        elseif r_track(j,i)+H > L
            box{i}(j,1:2) = [(L-2*H) L];
        else
            box{i}(j,1:2) = round([r_track(j,i)-H r_track(j,i)+H]);
        end
        if c_track(j,i)-H < 1
            box{i}(j,3:4) = [1 (2*H+1)];
        elseif c_track(j,i)+H > L
            box{i}(j,3:4) = [(L-2*H) L];
        else
            box{i}(j,3:4) = round([c_track(j,i)-H c_track(j,i)+H]);
        end
    end
end

%% Make little images around each cell!
for j = 1:nT
    for i = 1:nCells
        cinds = box{i}(j,1):box{i}(j,2);
        rinds = box{i}(j,3):box{i}(j,4);
        
        IM_cell{i}(:,:,j)  = IM(rinds,cinds,j);
    end
    fprintf('.')
end
fprintf('\n')
```
