## Supplementary Code 1 for "Engineering combinatorial and dynamic decoders using synthetic immediate-early genes": getCytoplasmicRegionCandidates.m

```
function BWc = getCytoplasmicRegionCandidates(Bt,Rt,BWt)
nT = size(BWt,3);

%%
% First, do a "test" segmentation of a few images to get centroids
cluster_intensities = [];
tv = round(linspace(1,nT,10));
for j = 1:length(tv)
    B  = Bt(:,:,tv(j));
    R  = Rt(:,:,tv(j));
    BW = BWt(:,:,tv(j));
    [TMP1 tmp2] = getCytRegion(R, B, BW, 15, 4);
    cluster_intensities = [cluster_intensities; tmp2];    
end

%%
% kmeans cluster this "test" set to get good initial guesses for
% cluster centroids
[idx mv] = kmeans(cluster_intensities, 4, 'emptyaction', 'singleton', 'replicates', 5);

% This also allows us to order the clusters as we want (in this case,
% sorted for increasing fluorescence)
[tmp I] = sort(sum(mv,2));
mv = mv(I,:);

%%
% Now re-cluster ALL the data using those centroids as initial guesses
cluster_intensities = [];
for j = 1:nT
    B  = Bt(:,:,j);
    R  = Rt(:,:,j);
    BW = BWt(:,:,j);
    [TMP1 tmp2] = getCytRegion(R, B, BW, 15, 4, mv);
    cluster_intensities = [cluster_intensities; tmp2];

    for k = 1:4
        BWc{1,k}(:,:,j) = imclose(TMP1(:,:,k), strel('disk', 2));
    end
end
```
