## Supplementary Code 1 for "Engineering combinatorial and dynamic decoders using synthetic immediate-early genes": getCytRegion.m

```
function [BWc mv] = getCytRegion(R, B, BWn, rdisk, nClusters, initialGuesses)
% function [BWc mv] = getCytRegion(R, B, BWn, rdisk, nClusters)
% 
% R     = red image
% B     = blue image
% BWn   = binary nuclear image
% rdisk = size of annulus to grow before "sculpting away" for nucleus
% nClus = # of distinct clusters for kmeans (should be 
% 
% BWc   = binary cytoplasmic images for each cluster
% mv    = the "mean values" - e.g. the cluster intensities

%% Grow an annulus around the nucleus - initial guess of cytoplasm
BWc = imdilate(BWn, strel('disk', rdisk)) & ~BWn;
inds = find(BWc);

%% Cluster pixel intensities
vals = zeros(length(inds),2);
vals(:,1) = double(R(inds));
vals(:,2) = double(B(inds));

% ik is the cluster membership for each nonzero pixel in BWc
if nargin < 6 || isempty(initialGuesses)
    [ik mv] = kmeans(vals, nClusters, 'emptyaction', 'singleton');
else
    [ik mv] = kmeans(vals, nClusters, 'emptyaction', 'singleton', 'start', initialGuesses);
end

%%
BW_cluster = zeros(size(BWn,1), size(BWn,2), nClusters);

for k = 1:nClusters
    % get an image of that cluster
    tmp = zeros(size(BWn));
    tmp(inds(ik == k)) = 1;
    tmp = imclose(tmp, strel('disk', 1));
    
	% Only keep one connected component of the cluster:
    % the biggest connected component after conjoining to the nucleus!
    CC = bwconncomp(tmp | BWn, 4);
    numPixels = cellfun(@numel,CC.PixelIdxList);
    [biggest,idx] = max(numPixels);
    
    tmp2 = zeros(size(tmp));
    tmp2(CC.PixelIdxList{idx}) = 1;
    tmp2 = imfill(tmp2, 'holes');
    tmp2 = tmp2 & ~BWn;
    
	BW_cluster(:,:,k) = tmp2;
end

BWc = BW_cluster;
```
