## Supplementary Code 1 for "Engineering combinatorial and dynamic decoders using synthetic immediate-early genes": getDistToBasin.m

```
function [d IM] = getDistToBasin(c1,c2,c3,maxval,x)
    
    IM = zeros(maxval(1),maxval(2));
    
    [m1 b1] = findLine(c1,c2);
    [m2 b2] = findLine(c3,c2);
    
    xx = linspace(1,maxval(1),1e3);
    y1 = m1*xx+b1;
    y2 = m2*xx+b2;
    
    for i = 1:length(xx)
        if round(y1(i)) > 0 & round(y1(i)) <= 255
            IM(round(xx(i)), round(y1(i))) = 1;
        end
        if round(y2(i)) > 0 & round(y1(i)) <= 255
            IM(round(xx(i)), round(y2(i))) = 1;
        end
    end
    
    yy = linspace(1,maxval(2),1e3);
    x1 = (yy-b1)/m1;
    x2 = (yy-b2)/m2;
    
    for i = 1:length(xx)
        if round(x1(i)) > 0 & round(x1(i)) <= 255
            IM(round(x1(i)), round(yy(i))) = 1;
        end
        if round(x2(i)) > 0 & round(x2(i)) <= 255
            IM(round(x2(i)), round(yy(i))) = 1;
        end
    end
    
    IM = IM > 0;
    
    IM = imfill(IM, round(c2)');
    IM = imopen(IM, strel('disk', 2));
    IM = bwdist(IM);
    
    % move 0-values to 1.
    x(x == 0) = 1;
    
    ind = sub2ind(double(maxval), double(x(:,1)), double(x(:,2)));
    
    d = IM(ind);
end

%%
function [m b] = findLine(l1,l2)
    % This helper function finds the line equidistant from 2 points
    
    m  = -(l2(1)-l1(1))/(l2(2)-l1(2));
    
    % a point on the line
    p = 1/2*[l1(1)+l2(1)
             l1(2)+l2(2)];
    
	b = p(2)-m*p(1);
end
```
