## Supplementary Code 1 for "Engineering combinatorial and dynamic decoders using synthetic immediate-early genes": getGoodCytoplasmicRegions.m

```
function OUT = getGoodCytoplasmicRegions(IN_k, isgood)

for i = 1:size(IN_k,1)
    OUT{i} = false(size(IN_k{i,1}));
    for j = 1:size(IN_k{i,1}, 3)
        tmp = false(size(IN_k{i,1}(:,:,1)));
        for k = 1:4
            if isgood(i,k)
                tmp = tmp | IN_k{i,k}(:,:,j);
            end
        end
        OUT{i}(:,:,j) = tmp;
    end
    fprintf('.')
end
fprintf('\n')
```
