## Supplementary Code 1 for "Engineering combinatorial and dynamic decoders using synthetic immediate-early genes": getNext.m

```
function next_index = getNext(T, current_index, current_frame)
% This function finds the next index corresponding to a tracked object
% given three inputs:
% 1 the current index
% 2 the current timepoint
% 3 the tracking object T (which maps the current to next)
%
% It returns either of 2 outputs:
% the next index IF SUCH AN INDEX EXISTS
% or, if the object is lost, it returns 0.

next_index = 0; % default return value

if ~current_index, return, end

v1 = T{current_frame}(:,1);
v2 = T{current_frame}(:,2);

i1 = find(v1 == current_index, 1, 'first');

% if no index, give up!
if isempty(i1), return, end

next_index = v2(i1);
```
