## Supplementary Code 1 for "Engineering combinatorial and dynamic decoders using synthetic immediate-early genes": getNucCytCorrection.m

```
function data = getNucCytCorrection(nuc, cyt, bkgd)

nT = length(nuc);

n = (nuc-bkgd);
c = (cyt-bkgd);
t = (1:nT)';

ii = ~isnan(n) & ~isnan(c);
t = t(ii);
n = n(ii);
c = c(ii);

Y = n;
X = [ones(length(t),1) c t];

coef = X \ Y;

% Plot a few things
x = c;
y = n;
yy = coef(1) + coef(2)*x + coef(3)*t;

data.erk_total     =  coef(1);
data.cyt_nuc_ratio = -coef(2);
data.kbleach       = -coef(3)/coef(1);
data.unbleach      = 1./diag(inv(diag(1-coef(3)/coef(1)*(t))));
data.nuc           = n;
data.cyt           = c;
data.nuc_corr      = n.*data.unbleach;
data.cyt_corr      = c.*data.unbleach;
data.nuc_tot_ratio = n/coef(1).*data.unbleach;
data.cyt_tot_ratio = c/coef(1)*-coef(2).*data.unbleach;
data.t             = t;

figure(1),clf
subplot(2,1,1)
plot(t,y,t,yy)
xlabel('time')
ylabel('nuc Erk')
legend({'raw nuclear' 'computed nuclear'});
subplot(2,1,2)
plot(data.t, data.nuc_tot_ratio, data.t, data.cyt_tot_ratio)
xlabel('time')
ylabel('fraction Erk')
```
