## Supplementary Code 1 for "Engineering combinatorial and dynamic decoders using synthetic immediate-early genes": getTrackedNuclei.m

```
function [Lout r_track c_track] = getTrackedNuclei(BW)
% function [Lout r_track c_track] = getTrackedNuclei(BW)
% 
% This function tracks objects through all frames based on finding the
% objects that "agree" on maximizing overlap by playing the movie forwards
% or backwards (i.e. that the next frame's object maximizes overlap with
% the previous frame, and that the previous frame maximizes overlap with
% the next frame). 
% 
% It takes the black & white movie of objects as input, and spits out
% a movie of only the tracked, labeled objects as well as two matrices,
% r_track and c_track, that contain their row & column centroid locations
% at each timepoint (and for each object).

nT = size(BW,3);

%% 1 - GET CENTROIDS OF ALL REGIONS
disp('(1/5) Getting centroids...')
clear rpc
for i = 1:nT
    rp = regionprops(BW(:,:,i), 'centroid');
    rpc{i} = reshape([rp.Centroid]',2,[])';
    fprintf('.')
end
fprintf('\n')

%% 2 - TRACK OBJECTS THROUGH ALL FRAMES
disp('(2/5) Tracking nuclei...')
for i = 1:nT-1
    T{i} = trackNuclei(BW(:,:,i),BW(:,:,i+1));
    fprintf('.')
end
fprintf('\n')

%% get about 80% of the data if you ask for perfection (not bad!)
%
% TO IMPLEMENT: finding objects that are "skipped" in one frame (by picking
% the intersected area of the frames on both sides)

disp('(3/5) get tracking matrix')
ir = zeros(size(T{1},1), length(T));
ir(:,1) = T{1}(:,1);

for j = 1:length(T)
    for i = 1:size(ir,1)
        ir(i,j+1) = getNext(T, ir(i,j), j);
    end
    fprintf('.')
end
fprintf('\n')

%%
disp('(4/5) making tracks...')

iperfect = find(all(ir > 0, 2));

r_track = zeros(nT,length(iperfect));
c_track = zeros(nT,length(iperfect));

for j = 1:nT
    for i = 1:length(iperfect)
        r_track(j,i) = rpc{j}(ir(iperfect(i),j),1);
        c_track(j,i) = rpc{j}(ir(iperfect(i),j),2);
    end
    fprintf('.')
end
fprintf('\n')

%% DEBUGGING: plot the tracks (to make sure tracking is working well)

% for i = 1:length(iperfect)
%     plot(r_track(:,i), c_track(:,i), 'o-')
%     set(gca, 'xlim', [0 1400], 'ylim', [0 1400])
%     title(i)
%     pause
% end

%% Get a final labeled object, BWout, that spans all timepoints
disp('(5/5) writing output image...')

Lout = zeros(size(BW));
for j = 1:nT
    L = bwlabel(BW(:,:,j));
    tmp = zeros(size(L));
    for i = 1:length(iperfect)
        ix = ir(iperfect(i),j);
        tmp(L == ix) = i;
    end
    Lout(:,:,j) = tmp;
    fprintf('.')
end
fprintf('\n')

disp('done!')
```
