## Supplementary Code 1 for "Engineering combinatorial and dynamic decoders using synthetic immediate-early genes": growCytRegion.m

```
function [BW mv] = growCytRegion(BWn, IM, rdisk, nClusters)
% INPUTS:
% BWn   = nuclear region
% IM    = fluorescent image of cell OR MULTIPLE IMAGES (i.e. BFP, YFP, RFP)
% rdisk = size of region to grow, before chipping away
% k     = number of kmeans clusters
% 
% OUTPUTS:
% BWc = cytoplasmic region

%% Grow a disk - this is the test cytoplasmic region
BWc = imdilate(BWn, strel('disk', rdisk)) & ~BWn;
inds = find(BWc);

%% Get n-dimensional intensities for each cyt pixel
vals = zeros(length(inds),size(IM,3));
for k = 1:size(IM,3)
    tmp = IM(:,:,k);
    vals(:,k) = tmp(inds);
end

%% Now kmeans cluster the data in vals
[ik mv] = kmeans(vals, nClusters, 'emptyaction', 'singleton');

%%
for k = 1:nClusters
    tmp = zeros(size(IM,1), size(IM,2));
    tmp(inds(ik == k)) = 1;
    tmp = imclose(tmp, strel('disk', 1));
    
    % HOW TO SELECT WHICH REGION TO KEEP?
    CC = bwconncomp(tmp | BWn);
    numPixels = cellfun(@numel,CC.PixelIdxList);
    [biggest,idx] = max(numPixels);
    tmp2 = zeros(size(tmp));
    tmp2(CC.PixelIdxList{idx}) = 1;
    tmp2 = tmp2 & ~BWn;
    
%     BW(:,:,k) = imclose(tmp2, strel('disk', 1));
    BW(:,:,k) = tmp2;
%     BW(:,:,k) = tmp;
end
% keyboard
return

%% Plot some results!
for i = 1:nClusters
%     tmp = zeros(size(IM,1), size(IM,2));
%     tmp(inds(ik == i)) = 1;
    tmp = BW(:,:,i);
    tmp = cat(3,tmp,cat(3,tmp,tmp));
    imshow(IM + 100*uint8(tmp))
%     drawnow
    pause(0.1)
    pause
end
% keyboard
```
