## Supplementary Code 1 for "Engineering combinatorial and dynamic decoders using synthetic immediate-early genes": imsmoothen.m

```
function BWout = imsmoothen(BW)
% nT = size(BW,3);
% BWout(:,:,1)  = BW(:,:,1);
% BWout(:,:,nT) = BW(:,:,nT);
% 
% for i = 2:nT-1
%     indsON  =  BW(:,:,i) | (sum(BW(:,:,i-1:i+1),3) > 2);
%     indsOFF = ~BW(:,:,i) | (sum(BW(:,:,i-1:i+1),3) < 2);
%     I = zeros(size(BW(:,:,i)));
%     I(indsON) = 1;
%     I(indsOFF) = 0;
%     BWout(:,:,i) = I;
% end

%%
nT = size(BW,3);
BWout = BW;

for i = 2:nT-1
    indsON  =  BWout(:,:,i) | (sum(BWout(:,:,i-1:i+1),3) > 1);
    I = zeros(size(BWout(:,:,i)));
    I(indsON) = 1;
    BWout(:,:,i) = I;
end
```
