## Supplementary Code 1 for "Engineering combinatorial and dynamic decoders using synthetic immediate-early genes": loadData.m

```
function [IM all_scales] = loadData(path, fname, iT, fix_scale)
% function IM = loadData(path, fname, iT, fix_scale)
% 
% This function loads the images from a .TIF stack (i.e. set of timepoints
% or z stacks) and converts the result to an 8 bit file. 
% 
% In order to fill out the dynamic range of the resulting 8 bit file (i.e.
% to use the values 0-255 relatively completely), it randomly samples
% 10,000 pixels for their intensity, and sets the 99th percentile intensity
% to a value of 127.
% 
% If the argument fix_scale is set to 1, it makes sure that every image in
% the TIF stack is scaled identically.
% 
% Input arguments:
% path      -   the path to the filename
% fname     -   the filename to load
% iT        -   the timepoints to load
% fix_scale -   keep the same scaling factor for all timepoints? (default
%               is zero)
% 
% Output arguments:
% IM        -   the output image stack
% 
% Example usage:
% IM_BFP  = loadData('DATA', 'overnight001c1.tif', 1:nT, 0);

%% Set default input argument
if nargin < 4 || isempty(fix_scale)
    fix_scale = 1;
end

%% Initialize some variables
scale = [];

% Initialize output image IM to the size: image rows x image columns x # of timepoints
tmp = imread(sprintf('%s/%s', path, fname), 1);
IM = uint8(zeros(size(tmp,1), size(tmp,2), length(iT)));

%% Load images

for i = 1:length(iT)
    % load image file
    tmp = imread(sprintf('%s/%s', path, fname), iT(i));
    
    % scale the loaded image and save to the ith element in IM
    [IM(:,:,i) scale] = scaleToUint8(tmp, scale);
    all_scales(i) = scale;
    % if not fix_scale, set scale to empty again
    if ~fix_scale
        scale = [];
    end
    fprintf('.')
end
fprintf('\n')

end % END MAIN FUNCTION


% DEFINE A HELPER FUNCTION, scaleToUint8
function [IM0 scale] = scaleToUint8(IM, scale)

% If you DON'T know the scaling factor, find it
if nargin < 2 || isempty(scale)
    
    % 1. randomly sample 10,000 pixels & get maximum in that set
    nSamples = 1e5;
    i_samp = round((numel(IM)-1)*rand(nSamples,1))+1;
    
    % measure intensities and sort in increasing order
    isort = sort(IM(i_samp));
    
    % the "max" intensity, around which we scale, is the 99% percentile
    maxim = isort(round(.99*nSamples));
    
    % convert to double for math
    maxim = double(maxim);
    
    SAFE_MAX = 127; % this is the value that we set the 99% percentile to.
    
    % thus, our scale factor is SAFE_MAX/maxim
    scale = (SAFE_MAX/maxim);
end

% now scale the image!
IM0 = uint8(round(double(IM)*scale));

end
```
