## Supplementary Code 1 for "Engineering combinatorial and dynamic decoders using synthetic immediate-early genes": measureAllIntensities.m

```
function i_out = measureAllIntensities(BW, IM, nClusters, icluster, closingRadius, erodeRadius, makeMovie, moviefile)
if nargin < 6 || isempty(erodeRadius)
    erodeRadius = 0;
end
if nargin < 7 || isempty(makeMovie)
    makeMovie = 0;
end
if nargin < 8 || isempty(moviefile)
    moviefile = 'tmp';
end

newMatlab = 1; % 1 if 2014; 0 if 2009

nCells = length(IM);
nT     = size(IM{1},3);
i_out  = nan*ones(nT, nCells);

warning off
for i = 1:nCells
    if makeMovie
        if newMatlab
            writerObj = VideoWriter(sprintf('%s_%0.3d.avi', moviefile, i), 'Motion JPEG AVI');
            open(writerObj);
        else
            aviobj = avifile(sprintf('%s_%0.3d.avi', moviefile, i), 'compression', 'none');
        end
    end
    for j = 1:nT
        I = IM{i}(:,:,j);
        B = BW{i}(:,:,j);
        if erodeRadius
            B = imerode(B, strel('disk', erodeRadius));
        end
        [i_out(j,i) IMout] = measureIntensity(B, I, nClusters, icluster, closingRadius);
        if isempty(IMout)
            IMout = false(size(I));
        end
        if makeMovie
            if newMatlab
                IMmovie = I+30*uint8(IMout);
                writeVideo(writerObj,IMmovie);
            else
                IMmovie = I+30*uint8(IMout);
                aviobj = addframe(aviobj, double(IMmovie)/255);
            end
        end
    end
    fprintf('.')
    if makeMovie
        if newMatlab
            close(writerObj);
        else
            aviobj = close(aviobj);
        end
    end
%     pause
end
fprintf('\n')
warning on
```
