## Supplementary Code 1 for "Engineering combinatorial and dynamic decoders using synthetic immediate-early genes": measureIntensity.m

```
function [intensity IMout] = measureIntensity(BW, IM, nClusters, icluster, closingRadius)
PLOT_ME = 1; % Use this for debugging, if you want to see what region you're measuring!

if nargin < 5 || isempty(closingRadius)
    closingRadius = 1;
end

inds = find(BW);
vals = double(IM(inds));

try
    if nClusters == 1
        % forget kmeans
        intensity = mean(vals);
    else
        % kmeans and then measure
        [idx mv] = kmeans(vals, nClusters, 'emptyaction', 'singleton', 'replicates', 3);
        [mv I] = sort(mv);
        
        tmp = zeros(size(IM));
        tmp(inds(idx == I(icluster))) = 1;
        tmp = tmp > 0;
        tmp = imclose(tmp, strel('disk', closingRadius));
        
        if PLOT_ME
            imshow(IM+30*uint8(tmp),[0 150])
            drawnow
        end
        
%         intensity = mean(vals(idx == I(icluster)));
        intensity = mean(double(IM(tmp)));
        IMout = tmp;
    end
catch
    intensity = nan;
    IMout = [];
end
```
