## Supplementary Code 1 for "Engineering combinatorial and dynamic decoders using synthetic immediate-early genes": oldOrderClusters.m

```
function k_xy = orderClusters(coords, nClusters)

[idx cents] = kmeans(coords, nClusters, 'emptyaction', 'singleton', 'replicates', 5);

for k = 1:nClusters
    plot(coords(:,1), coords(:,2), '.', ...
         coords(idx == k,1), coords(idx == k,2), 'o')
     pause
end

% now for each, assign which one is closest
% i can at least brute force this
nT = size(coords,1)/4;

% %%
% for i = 1:nT
%     ii = (1:4)+(i-1)*4;
%     plot(coords(:,1), coords(:,2), '.', coords(ii,1), coords(ii,2), 'o')
%     pause
% end

%% OK, for each point, find the cluster it's closest to.
for i = 1:nT
    ii = (1:4)+(i-1)*4;
    cur_point = coords(ii,:);
    plot(coords(:,1), coords(:,2), '.', cur_point(:,1), cur_point(:,2), 'o')
    k_xy(i,:) = getMutualMatches(cur_point, cents)';
end

end

function k_xy = getMutualMatches(xy, xy0)
n = size(xy,1);
best1 = zeros(n,1);
best2 = zeros(n,1);
k_xy  = nan*ones(n,1);

for i = 1:n
    d1 = (xy(i,1) - xy0(:,1)).^2 + (xy(i,2) - xy0(:,2)).^2;
    best1(i) = find(d1 == min(d1), 1, 'first');
    d2 = (xy(:,1) - xy0(i,1)).^2 + (xy(:,2) - xy0(i,2)).^2;
    best2(i) = find(d2 == min(d2), 1, 'first');
end

k_xy(best2(best1) == (1:n)') = best1(best2(best1) == (1:n)');

end
```
