## Supplementary Code 1 for "Engineering combinatorial and dynamic decoders using synthetic immediate-early genes": thresholdImage2.m

```
function BW = thresholdImage2(IM, nClusters)

% Subsample image for kmeans
A_IM = size(IM,1)*size(IM,2);
nSamples = min(A_IM, 1e4);
i_samp = ceil(A_IM*rand(nSamples,1));

for i = 1:size(IM,3)
    tmp = IM(:,:,i);
    ivals(:,i) = double(tmp(i_samp));
end

[idx c] = kmeans(ivals, nClusters, 'emptyaction', 'singleton', 'start', [20 20; 20 200; 200 20]);

%% Get distance of ALL points to 3 clusters

for i = 1:nClusters
    dist(:,:,i) = (double(IM(:,:,1))-c(i,1)).^2 + ...
                  (double(IM(:,:,2))-c(i,2)).^2;
end

%% You can also plot the points & what cluster they are in
%  This is commented out because it doesn't need be done once this has been
%  validated!

% for i = 1:nClusters
%     plot(ivals(       :,1), ivals(       :,2), '.', ...
%          ivals(idx == i,1), ivals(idx == i,2), 'o')
%     pause
% end

%% MUCH more useful: DISTANCE in kmeans space
% Now that you have this, you can get the distance from a kmeans cluster
% (low RFP, high irFP) and find only those pixels that are CLOSE to this
% centroid!

D = dist(:,:,2);
% D = D-imtophat(D, strel('ball', 4, 1e4));
D = D-imtophat(D, strel('disk', 15));

%% Subsample this distance image to get the threshold for being "close"
Ds = log10(D(i_samp));
imshow(D,[])
[idx2 c] = kmeans(Ds, 3, 'emptyaction', 'singleton', 'start', [3 3.5 4]');
c = 10.^sort(c);

%% This has its own distance image

for i = 1:3
    dist2(:,:,i) = (D-c(i)).^2;
end

%% An image of which cluster you're closest to

I = zeros(size(IM,1),size(IM,2));
mindist = min(dist2,[],3);
for i = 1:nClusters
    I(dist2(:,:,i) == mindist) = i;
end

% keyboard

%% Now keep those that are members of cluster 1!
BW = I == 1 | I == 2;

BW = imopen(BW, strel('disk', 1));
BW = bwfill(BW,'holes');
% imshow(BW)
```
