## Supplementary Code 1 for "Engineering combinatorial and dynamic decoders using synthetic immediate-early genes": thresholdImage.m

```
function BW = thresholdImage(IM, thr, above)
warning off

BW = true(size(IM,1), size(IM,2));

for i = 1:size(IM,3)    
    TMP = IM(:,:,i);
    
    if above(i)
        BW = BW & TMP > thr(i);
    else
        BW = BW & TMP < thr(i);
    end
end

BW = BW-imtophat(BW, strel('disk', 9));

warning on
```
