## Supplementary Code 1 for "Engineering combinatorial and dynamic decoders using synthetic immediate-early genes": thresholdImage_Phy_BFP.m

```
function BW = thresholdImage_Phy_Nuc(R, B, dist_thr)
% function BW = thresholdImage_Phy_Nuc(R, B)
% 
% This function takes in two images, a red Phy and infrared nuclear image,
% as well as the number of "clusters" that represent different types of
% cellular features. We typically pick 3 clusters representing:
% CELL NUCLEI       - these should have bright B and dim R
% CELL CYTOPLASM    - these should have brighter red and dim B
% NON-CELL BKGD     - these should be dim in both channels


%% INITIALIZE SOME VARIABLES
% the # of clusters and initial guesses for each cluster's intensity
initialguess = [20  20
                20  80
                100 10
                50  50
                30  70];

nClusters = size(initialguess,1);

% The # of pixels in the images
A_IM = numel(R);

%% Do kmeans clustering
% Subsample image for kmeans
nSamples = min(A_IM, 1e4); % sample this many pixels
i_samp = round((A_IM-1)*rand(nSamples,1))+1;

% Get pixel intensities and save into variable "ivals"
ivals(:,1) = double(B(i_samp));
ivals(:,2) = double(R(i_samp));

% This is where we do the clustering! The important output is "c" - the
% cluster centers.
[idx c] = kmeans(ivals, nClusters, 'emptyaction', 'singleton', 'start', initialguess);

D = getDistToBasin(c(5,:)', c(3,:)', c(2,:)', [255 255], [B(:) R(:)]);
D = reshape(D, size(R));
D = sqrt(D);

%%
D = D-imtophat(D, strel('disk', 15));

BW = D < dist_thr;

% Postprocess by removing tiny objects and filling holes
BW = bwfill(logical(BW),'holes');
BW = imopen(logical(BW), strel('disk', 7));
```
