## Supplementary Code 1 for "Engineering combinatorial and dynamic decoders using synthetic immediate-early genes": trackNuclei.m

```
function T12 = trackNuclei(IM1, IM2)

    [L1 n1] = bwlabel(IM1);
    [L2 n2] = bwlabel(IM2);
    
    % Get forward hits
    h12 = getHits(L1,n1,L2);
    
    % Get reverse hits
    h21 = getHits(L2,n2,L1);
    
    % Figure out "bidirectional" hits - where h12 and h21 point to each
    % other
    i12 = find(h12 > 0);
    v2 = h12(i12(h12(i12) == h12(h21(h12(i12)))));
    v1 = h21(v2);
    
    T12 = [v1 v2];
end

function h1 = getHits(L1,n1,L2)
    h1 = zeros(n1,1);
    for i = 1:n1
        inds = find(L1 == i & ...  % ith region in L1 AND
                    L2 > 0);       % ... nonzero in L2
        
        if isempty(inds), continue, end % if no overlap, skip!
        
        h1(i) = mode(L2(inds)); % get the mode - highest overlap
    end
end
```
