## Supplementary figures and images for "Engineering combinatorial and dynamic decoders using synthetic immediate-early genes"

### V_BandPass001-MaxIP.nd2 - Z=5 C=0.tif

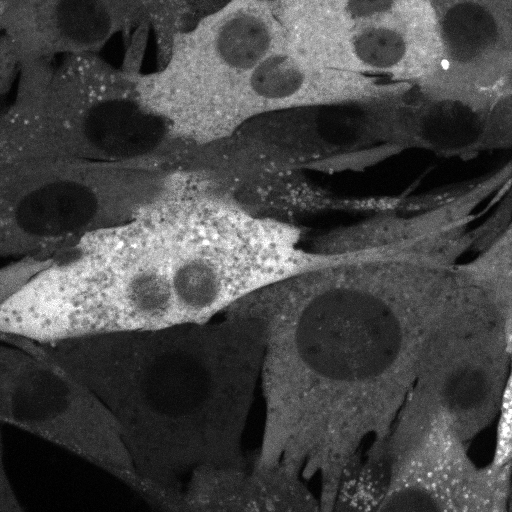
