## Supplementary Code 2 for "Engineering combinatorial and dynamic decoders using synthetic immediate-early genes": Fluorescence Analysis Code.R

```
install.packages('ggplot2')
install.packages('data.table')
install.packages('plotrix')
library(ggplot2)
library(data.table)
library(plotrix)

#Function to read text files and puts them into a large data frame
#arguments are (root); (file) folder path variables, timestep - how much time between, 
#(gene) name of cells
dfYFP<-function(root,files,timestep,gene){
  d <- data.frame()
  
  for(i in 1:length(files)){
    tmp <- files[i]
    tmpSplit <- strsplit(tmp, '-')
    dtmp<- read.csv(paste(c(root, files[i]), collapse = ''),header = T, sep='\t')
    if("Frame" %in% colnames(dtmp)) {
      dtmp<- subset (dtmp, select = -(Frame))
    }
    #if (dtmp$Slice[1] != 1) {next}
    
    #Add factors for identification
    dftmp <- as.data.frame(dtmp)
    dftmp$Position <- rep(tmpSplit[[1]][1], nrow(dftmp))
    dftmp$IV <- rep(tmpSplit[[1]][2], nrow(dftmp))
    if (dftmp$IV == "Combo"){
      dftmp$IV = "DoxSerum"
    }
    dftmp$Cell <- rep(tmpSplit[[1]][3], nrow(dftmp))
    dftmp$Gene <- rep(gene, nrow(dftmp))
    slices <- dftmp$Slice

    
    for (j in 1:length(slices)) {
      dftmp$ID[j] <- rep (paste(c(tmpSplit[[1]][1],tmpSplit[[1]][2],slices[j]), collapse = ''))
      dftmp$ID2[j] <- rep (paste(c(tmpSplit[[1]][2],slices[j]), collapse = ''))
      dftmp$ID3[j] <- rep(paste(c(tmp)))   
    }
    
    dftmp$Time <- dftmp$Slice * timestep - timestep
    if (i==1) {
      df = dftmp
    }
    else df <- rbind(df, dftmp)
  } 
  return (df)
}

#Function takes data frame and calculates statistics 
#arguments: (df) dataframe from dfYFP, (sliceskip) how many timepoints to skip before analysis
# (bkgd) value of background intensity and (normtech) 0, 1, 2 => each a different normalization technique
dtYFP<- function (df,sliceskip,bkgd2, normtech){
  z <- data.table(df)
  z <- subset(z, Slice >= sliceskip)
  #Background normalization
  background <- subset(z,Cell == "cell_bkgd.txt")
  data.noback <- subset(z, Cell != "cell_bkgd.txt")
  data.noback$bkgd = background$Mean[chmatch(as.character(data.noback$ID), as.character(background$ID))]
  data.noback$bsub <-  data.noback$Mean/data.noback$bkgd - 1
  z <- data.noback
  z[, min := mean(head(Mean , 4)), by = c("Position" , "IV", "Cell")]
  z[, bmin := mean(head(bsub , 4)), by = c("Position" , "IV", "Cell")]
  z[, max := mean(tail(Mean , 4)), by = c("Position" , "IV", "Cell")]
  if (normtech == 0) {
    z[, normmean := (Mean-min), by = c("ID", "Slice", "Cell")]  #Change in FU Normalization
  }
  if (normtech == 1) {
    z[, normmean := (Mean-bkgd2)/(min-bkgd2), by = c("ID", "Slice", "Cell")]  #Fold Change
  }  
  if (normtech == 2) {
    z[, normmean := (Mean-bkgd2), by = c("ID", "Slice", "Cell")]  #Raw Data
  }
  if (normtech == 3) {
    z[, normmean := (bsub-bmin), by = c("ID", "Slice", "Cell")]  #Background subtraction 
  }
  z[, mean := mean(normmean,na.rm = TRUE), by = c("IV", "Slice")]
  z[, stddev := sd(normmean, na.rm = TRUE), by = c("IV", "Slice")]
  z[, n := length(ID2), by = c("IV", "Slice")]
  z[, se := stddev/sqrt(n), by = c("IV", "Slice")]
  return (z)
}

#takes datatable from dtYFP and plots each cell individually
dfplot<-function(z){
  ggplot(z, aes(x=Time, y=normmean, color=ID3)) + geom_line() +
    facet_wrap(Position~IV) 
}

#takes datatable from dtYFP and plots mean+/-sem for each condition
dtplot<-function(z){
  p <- ggplot(z, aes(x=Time, y=mean, color = IV)) + geom_line() +
    facet_wrap(~Gene) 
  p <- p + labs(x = "Time (min)", y = ("Min Normalized Mean Intensity")) + 
    geom_ribbon(aes(ymin = mean-se, ymax=mean+se), linetype=2, alpha=0.1) 
  p
}

#datatable from dtYFP and plots individual trajectories overlayed with mean trajectory
#broken down by condition
dfdtplot <- function(z){
  p <- ggplot(z, aes(x=Time, y = mean))  + 
    geom_line(data = z, aes(x=Time, y = normmean, Group = ID3), color = "grey") + 
    geom_line(size=1, alpha=0.8) +
    facet_wrap(~IV)
  p
}

#Creates a Bar Graph, graphing foldchange over baseline
barplot2 <- function(z) {
  z <- unique(z, by=c("mean"))  
  z[,baseline := mean(head(mean, 1)) , by = "IV"]
  z$foldchange = z$mean
  z <- subset(z, z$Slice == tail(z$Slice,1))
  z$IV <- factor(z$IV,levels = c("Starve", "Serum", "Dox", "DoxSerum"))
  
  ggplot(data=z, aes(x=IV, y=foldchange)) +
    geom_bar(stat="identity", position=position_dodge()) +
    geom_errorbar(aes(ymin=foldchange-se, ymax=foldchange+se), width=.2,
                  position=position_dodge(.9))
}

# Function to produce summary statistics (mean and +/- sd)
data_summary <- function(x) {
  m <- mean(x)
  ymin <- m-sd(x)
  ymax <- m+sd(x)
  return(c(y=m,ymin=ymin,ymax=ymax))
}


#taken from https://rdrr.io/cran/MESS/src/R/auc.R
#calculates area under the curve
auc <-
  function(x, y, from = min(x, na.rm=TRUE), to = max(x, na.rm=TRUE), type=c("linear", "spline"), absolutearea=FALSE, ...)
  {
    type <- match.arg(type)
    
    # Sanity checks
    stopifnot(length(x) == length(y))
    
    stopifnot(!is.na(from))
    
    if (length(unique(x)) < 2)
      return(NA)
    
    if (type=="linear") {
      
      ## Default option
      if (absolutearea==FALSE) {
        values <- approx(x, y, xout = sort(unique(c(from, to, x[x > from & x < to]))), ...)
        res <- 0.5 * sum(diff(values$x) * (values$y[-1] + values$y[-length(values$y)]))
      } else { ## Absolute areas
        ## This is done by adding artificial dummy points on the x axis
        o <- order(x)
        ox <- x[o]
        oy <- y[o]
        
        idx <- which(diff(oy >= 0)!=0)
        newx <- c(x, x[idx] - oy[idx]*(x[idx+1]-x[idx]) / (y[idx+1]-y[idx]))
        newy <- c(y, rep(0, length(idx)))
        values <- approx(newx, newy, xout = sort(unique(c(from, to, newx[newx > from & newx < to]))), ...)
        res <- 0.5 * sum(diff(values$x) * (abs(values$y[-1]) + abs(values$y[-length(values$y)])))
      }
      
    } else { ## If it is not linear approximation
      if (absolutearea)
        myfunction <- function(z) { abs(splinefun(x, y, method="natural")(z)) }
      else
        myfunction <- splinefun(x, y, method="natural")
      
      res <- integrate(myfunction, lower=from, upper=to)$value
    }
    
    res
  }

#takes the datatable from dtYFP and argument (bkgd) value of background 
#gives a datable that includes the area under the curve of each trajectory
dtauc <- function (dt,bkgd) {  
  dt[,bkgdsub:= Mean - bkgd]
  dt[, newmin:= mean(head(bkgdsub , 5)), by = c("Position" , "IV", "Cell")]
  dt[, data:= bkgdsub - newmin]
  dt[, AUC:= auc(Time,data), by="ID3"]
  dt[, meanAUC:= mean(AUC) , by="IV"]
  dt[, stdAUC:= sd(AUC), by = "IV"]
  return (dt)
}

# plots out a dotplot of the auc for each condition 
aucplot <- function(z) {
  z <- unique(z, by= c("AUC"))
  p<-ggplot(z, aes(x=IV, y=AUC)) + 
    geom_dotplot(binaxis='y', stackdir='center', dotsize= 0.5)
  p <- p + stat_summary(fun.data=data_summary, color="red", size = 0.5)
  p <- p + scale_x_discrete(limits=c("Starve", "Dox", "Serum1", "DoxSerum1"))
  p
}

#Example
#Data for Figure S4D
files<- list.files ("/Users/Pavithran/Desktop/Research/Toettcher_Lab/Image_analysis/Ravindran et al Paper/Sample Data Set/")
root<- "/Users/Pavithran/Desktop/Research/Toettcher_Lab/Image_analysis/Ravindran et al Paper/Sample Data Set/"
df = dfYFP(root,files,5,"Clones")
dt = dtYFP(df,1,1353,1)
dtplot(dt)
dfdtplot(dt)
```
